# Generalized additive models to analyze non-linear trends in biomedical longitudinal data using R: Beyond repeated measures ANOVA and Linear Mixed Models

**DOI:** 10.1101/2021.06.10.447970

**Authors:** Ariel I. Mundo, Timothy J. Muldoon, John R. Tipton

**Affiliations:** Department of Biomedical Engineering, University of Arkansas, Fayetteville, AR, USA; Department of Mathematical Sciences, University of Arkansas, Fayetteville, AR, USA

**Keywords:** longitudinal data, biomedical data, generalized additive models, simulation, R

## Abstract

In biomedical research, the outcome of longitudinal studies has been traditionally analyzed using the *repeated measures analysis of variance* (rm-ANOVA) or more recently, *linear mixed models* (LMEMs). Although LMEMs are less restrictive than rm-ANOVA as they can work with unbalanced data and non-constant correlation between observations, both methodologies assume a linear trend in the measured response. It is common in biomedical research that the true trend response is nonlinear and in these cases the linearity assumption of rm-ANOVA and LMEMs can lead to biased estimates and unreliable inference.

In contrast, GAMs relax the linearity assumption of rm-ANOVA and LMEMs and allow the data to determine the fit of the model while also permitting incomplete observations and different correlation structures. Therefore, GAMs present an excellent choice to analyze longitudinal data with non-linear trends in the context of biomedical research. This paper summarizes the limitations of rm-ANOVA and LMEMs and uses simulated data to visually show how both methods produce biased estimates when used on data with non-linear trends. We present the basic theory of GAMs and using reported trends of oxygen saturation in tumors, we simulate example longitudinal data (2 treatment groups, 10 subjects per group, 5 repeated measures for each group) to demonstrate their implementation in R. We also show that GAMs are able to produce estimates with non-linear trends even when incomplete observations exist (with 40% of the simulated observations missing). To make this work reproducible, the code and data used in this paper are available at: https://github.com/aimundo/GAMs-biomedical-research.

## 2 Background

Longitudinal studies are designed to repeatedly measure a variable of interest in a group (or groups) of subjects, with the intention of observing the evolution of effect across time rather than analyzing a single time point (e.g., a crosssectional study). Biomedical research frequently uses longitudinal studies to analyze the evolution of a “treatment” effect across multiple time points, with subjects of analysis ranging from animals (mice, rats, rabbits), to human patients, cells, or blood samples, among many others. Tumor response^1–4^, antibody expression^5,6^, and cell metabolism^7,8^ are examples of different situations where researchers have used longitudinal designs to study some physiological response. Because the frequency of the measurements in a longitudinal study is dependent on the biological phenomena of interest and the experimental design of the study, the frequency of such measurements can range from minute intervals to study a short-term response such as anesthesia effects in animals^9^, to weekly measurements to analyze a mid-term response like the evolution of dermatitis symptoms in breast cancer patients^10^, to monthly measurements to study a long-term response such as mouth opening following radiotherapy (RT) in neck cancer patients^11^.

Traditionally, a “frequentist” or “classical” statistical paradigm is used in biomedical research to derive inferences from a longitudinal study. The frequentist paradigm regards probability as the limit of the expected outcome when an experiment is repeated a large number of times^12^, and such view is applied to the analysis of longitudinal data by assuming a null hypothesis under a statistical model that is often an *analysis of variance over repeated measures* (repeated measures ANOVA or rm-ANOVA). The rm-ANOVA model makes three assumptions regarding longitudinal data: 1) linearity of the response across time, 2) constant correlation across same-subject measurements, and 3) observations from each subject are obtained at all time points through the study (a condition also known as *complete observations*)^13,14^.

The expected linear behavior of the response through time is a key requisite in rm-ANOVA^15^. This “linearity assumption” in rm-ANOVA implies that the model is misspecified when the data does not follow a linear trend, which results in unreliable inference. In longitudinal biomedical research, non-linear trends are the norm rather than the exception. A particular example of this non-linear behavior in longitudinal data arises in measurements of tumor response to chemo and/or radiotherapy in preclinical and clinical settings^1,8,16^. These studies have shown that the collected signal does not follow a linear trend over time, and presents extreme variability at different time points, making the fit of rm-ANOVA model inconsistent with the observed variation. Therefore, when rm-ANOVA is used to draw inference of such data the estimates are inevitably biased because the model is only able to accommodate linear trends that fail to adequately represent the biological phenomenon of interest.

A *post hoc* analysis is often used in conjunction with rm-ANOVA to perform repeated comparisons to estimate a *p-value*, which in turn is used as a measure of significance. Although it is possible that a *post hoc* analysis of rm-ANOVA is able to find “significant effects (*p-value*<0.05)” from data with non-linear trends, the validity of such a metric is dependent on how adequate the model fits the data. In other words, *p-values* are valid only if the model and the data have good agreement; if that is not the case, a “Type III” error (known as “model misspecification”) occurs^17^. For example, model misspecification will occur when a model that is only able to explain linear responses (such as rm-ANOVA) is fitted to data that follows a quadratic trend, thereby causing the resulting *p-values* and parameter estimates to be invalid^18^.

Additionally, the *p-value* itself is highly variable, and multiple comparisons can inflate the false positivity rate (Type I error or *α*)^19,20^, consequently biasing the conclusions of the study. Corrections exist to address the Type I error issue of multiple comparisons (such as Bonferroni^21^), but they in turn reduce statistical power (1-*β*)^22^, and lead to increased Type II error (failing to reject the null hypothesis when it is false)^23,24^. Therefore, the tradeoff of *post hoc* comparisons in rm-ANOVA between Type I, II and III errors might be difficult to resolve in a biomedical longitudinal study where a delicate balance exists between statistical power and sample size.

On the other hand, the assumption of constant correlation in rm-ANOVA (often known as the *compound symmetry assumption*) is typically unreasonable because correlation between the measured responses often diminishes as the time interval between the observation increases^25^. Corrections can be made in rm-ANOVA in the absence of compound symmetry^26,27^, but the effectiveness of the correction is limited by the size of the sample, the number of measurements^28^, and group sizes^29^. In the case of biomedical research, where living subjects are frequently used, sample sizes are often not “large” due to ethical and budgetary reasons,^30^ which might cause the corrections for lack of compound symmetry to be ineffective.

Due to a variety of causes, the number of observations during a study can be different between all subjects. For example, in a clinical trial patients may voluntarily withdraw, whereas attrition due to injury or weight loss in preclinical animal studies is possible. It is even plausible that unexpected complications with equipment or supplies arise that prevent the researcher from collecting measurements at certain time points. In each of these scenarios, the *complete observations* assumption of classical rm-ANOVA is violated. When incomplete observations occur, a rm-ANOVA model is fit by excluding all subjects with incomplete observations from the analysis^13^. This elimination of partially missing data from the analysis can result in increased costs if the desired statistical power is not met with the remaining observations, because it would be necessary to enroll more subjects. At the same time, if the excluded observations contain insightful information that is not used, their elimination from the analysis may limit the demonstration of significant differences between groups.

During the last decade, the biomedical community has started to recognize the limitations of rm-ANOVA in the analysis of longitudinal data. The recognition on the shortcomings of rm-ANOVA is exemplified by the use of linear mixed effects models (LMEMs) by certain groups to analyze longitudinal tumor response data^8,16^. Briefly, LMEMs incorporate *fixed effects*, which correspond to the levels of experimental factors in the study (e.g., the different drug regimens in a clinical trial), and *random effects*, which account for random variation within the population (e.g., the individual-level differences not due to treatment such as weight or age). When compared to the traditional rm-ANOVA, LMEMs are more flexible as they can accommodate incomplete observations for multiple subjects and allow different modeling strategies for the variability within each measure in every subject^15^. However, LMEMs impose restrictions in the distribution of the random effects, which need to be independent^13,31^. And even more importantly, LMEMs also assume by default a linear relationship between the response and time^15^ (polynomial effects can be used, but this approach has its own shortcomings as we discuss in Section 4.2.1).

As the rm-ANOVA and the more flexible LMEM approaches make overly restrictive assumptions regarding the trend of the response, there is a need for biomedical researchers to explore the use of statistical tools that allow the data (and not a model assumption) to determine the trend of the fitted model and to enable appropriate inference. In this regard, generalized additive models (GAMs) present an alternative approach to analyze longitudinal data. Although not frequently used by the biomedical community, these semi-parametric models are customarily used in other fields to analyze longitudinal data. Examples of the use of GAMs include the analysis of temporal variations in geochemical and palaeoecological data^32–34^, health-environment interactions^35^ and the dynamics of government in political science^36^. There are several advantages of GAMs over LMEMs and rm-ANOVA models: 1) GAMs can fit a more flexible class of smooth responses that enable the data to dictate the trend in the fit of the model, 2) they can model non-constant correlation between repeated measurements^37^, and 3) can easily accommodate incomplete observations. Therefore, GAMs provide a more flexible statistical approach to analyze non-linear biomedical longitudinal data than LMEMs and rm-ANOVA.

The current advances in programming languages designed for statistical analysis (specifically R), have eased the computational implementation of traditional models such as rm-ANOVA and more complex approaches such as LMEMs and GAMs. In particular, R^38^ has an extensive collection of documentation and functions to fit GAMs in the package *mgcv*^37,39^ that speed up the initial stages of the analysis and enable the use of advanced modeling structures (e.g. hierarchical models, confidence interval comparisons) without requiring advanced programming skills. At the same time, R has many tools that simplify data simulation.

Data simulation methods are an emerging technique that allow the researcher to create and explore different alternatives for analysis without collecting information in the field, reducing the time window between experiment design and its implementation. In addition, simulation can be also used for power calculations and study design questions^28^.

This work provides biomedical researchers with a clear understanding of the theory and the practice of using GAMs to analyze longitudinal data using by focusing on four areas. First, the limitations of LMEMs and rm-ANOVA regarding an expected trend of the response, constant correlation structures, and complete observations are explained in detail. Second, the key theoretical elements of GAMs are presented using clear and simple mathematical notation while explaining the context and interpretation of the equations. Third, we illustrate the type of non-linear longitudinal data that often occurs in biomedical research using simulated data that reproduces patterns in previously reported studies^16^. The simulated data experiments highlight the differences in inference between rm-ANOVA, LMEMs and GAMs on data similar to what is commonly observed in biomedical studies. Finally, reproducibility is emphasized by providing the code to generate the simulated data and the implementation of different models in R, in conjunction with a step-by-step guide demonstrating how to fit models of increasing complexity.

In summary, this work will allow biomedical researchers to identify when the use of GAMs instead of rm-ANOVA or LMEMs is appropriate to analyze longitudinal data, and provide guidance on the implementation of these models to improve the standards for reproducibility in biomedical research.

## 3 Challenges presented by longitudinal studies

### 3.1 The repeated measures ANOVA and Linear Mixed Model

The *repeated measures analysis of variance* (rm-ANOVA) and the *linear mixed model* (LMEM) are the most commonly used statistical analysis for longitudinal data in biomedical research. These statistical methodologies require certain assumptions for the model to be valid. From a practical view, the assumptions can be divided in three areas: 1) an assumed relationship between covariates and response, 2) a constant correlation between measurements, and, 3) complete observations for all subjects. Each one of these assumptions is discussed below.

### 3.2 Assumed relationship

#### 3.2.1 The repeated measures ANOVA case

In a longitudinal biomedical study, two or more groups of subjects (e.g., human subject, mice, samples) are subject to different treatments (e.g., a “treatment” group receives a novel drug or intervention vs. a “control” group that receives a placebo), and measurements from each subject within each group are collected at specific time points. The collected response is modeled with *fixed* components. The *fixed* component can be understood as a constant value in the response which the researcher is interested in measuring, i.e., the average effect of the novel drug/intervention in the “treatment” group.

Mathematically speaking, a rm-ANOVA model with an interaction can be written as

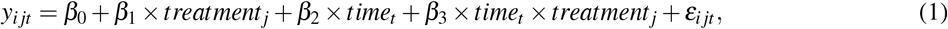

In this model *y*_*i jt*_ is the response for subject *i*, in treatment group *j* at time *t*, which can be decomposed in a mean value *β*_0_, *fixed effects* of treatment (*treatment*_*j*_), time (*time*_*t*_), and their interaction *time*_*t*_ *×treatment*_*j*_ which have linear slopes given by *β*_1_, *β*_2_ and *β*_3_, respectively. Independent errors *ε*_*i jt*_ represent random variation from the sampling process assumed to be 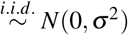 (independently and identically normally distributed with mean zero and variance *σ* ^2^). In a biomedical research context, suppose two treatments groups are used in a study (e.g., “placebo” vs. “novel drug”, or “saline” vs. “chemotherapy”). Then, the group terms in Equation (1) can be written as below with *treatment*_*j*_ = 0 representing the first treatment group (Group A) and *treatment*_*j*_ = 1 representing the second treatment group (Group B). With this notation, the linear model then can be expressed as

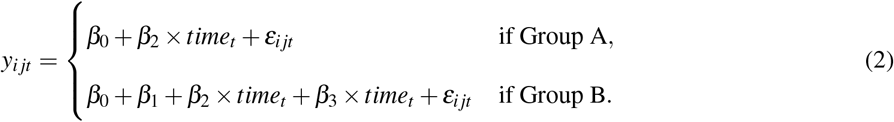

To further simplify the expression, substitute 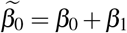 and 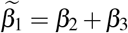 in the equation for Group B. This substitution allows for a different intercept and slope for Groups A and B. The model is then written as

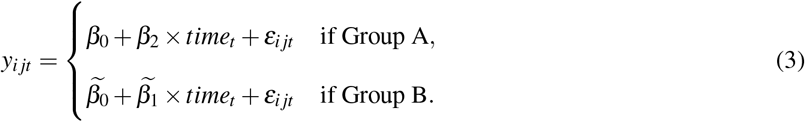

Presenting the model in this manner makes clear that when treating different groups, an rm-ANOVA model is able to accommodate non-parallel lines in each case (different intercepts and slopes per group). In other words, the rm-ANOVA model “expects” a linear relationship between the covariates and the response. This means that either presented as Equations (1), (2) or (3), an rm-ANOVA model is only able to accommodate linear patterns in the data. If the data show non-linear trends, the rm-ANOVA model will approximate this behavior with non-parallel lines.

#### 3.2.2 The Linear Mixed Model (LMEM) Case

A LMEM is a class of statistical models that incorporates *fixed effects* to model the relationship between the covariates and the response, and *random effects* to model subject variability that is not the primary focus of the study but that might be important to account for^15,40^. A LMEM with interaction between time and treatment for a longitudinal study can be written as

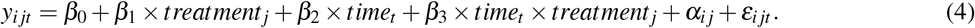

When Equations (1) and (4) are compared, it is noticeable that LMEMs and rm-ANOVA have the same construction regarding the *fixed effects* of time and treatment, but that the LMEM incorporates an additional source of variation (the term *α*_*i j*_). This term *α*_*i j*_ corresponds to the *random effect*, accounting for variability in each subject (subject_*i*_) within each group (group _*j*_). The *random* component can also be understood as modeling some “noise” in the response, but that does not arise from the sampling error term *ε*_*i jt*_ from Equations (1) through (3).

For example, if the blood concentration of a drug is measured in certain subjects in the early hours of the morning while other subjects are measured in the afternoon, it is possible that the difference in the collection time introduces some “noise” in the data that needs to be accounted for. As the name suggests, this “random” variability needs to be modeled as a variable rather than as a constant value. The random effect *α*_*i j*_ in Equation (4) is assumed to be 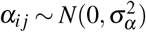. In essence, the *random effect* in a LMEM enables fitting models with different intercepts at the subject-level^15^. However, the expected linear relationship of the covariates and the response in Equation (1) and in Equation (4) is essentially the same, representing a major limitation of LMEMs to fit a non-linear response.

Of note, LMEMs are capable of fitting non-linear trends using an “empirical” approach (using polynomial fixed effects instead of linear effects such as in Equation (4)), which is described in detail by Pinheiro and Bates^15^. However, polynomial fits have limited predictive power, cause bias on the boundaries of the covariates^36^, and more importantly, their lack of biological or mechanistic interpretation limits their use in biomedical studies^15^.

### 3.3 Covariance in rm-ANOVA and LMEMs

In a longitudinal study there is an expected *covariance* between repeated measurements on the same subject, and because repeated measures occur in the subjects within each group, there is a *covariance* between measurements at each time point within each group. The *covariance matrix* (also known as the variance-covariance matrix) is a matrix that captures the variation between and within subjects in a longitudinal study^41^ (For an in-depth analysis of the covariance matrix see West^40^ and Weiss^42^).

In the case of an rm-ANOVA analysis, it is typically assumed that the covariance matrix has a specific construction known as *compound symmetry* (also known as “sphericity” or “circularity”). Under this assumption, the between-subject variance and within-subject correlation are constant across time^26,42,43^. However, it has been shown that this condition is frequently not justified because the correlation between measurements tends to change over time^44^; and is higher between consecutive measurements^13,25^. Although corrections can be made (such as Huyhn-Feldt or Greenhouse-Geisser)^26,27^ their effectiveness is dependent on sample size and number of repeated measurements^28^, and it has been shown that corrections are not robust if the group sizes are unbalanced^29^. Because biomedical longitudinal studies are often limited in sample size and can have an imbalanced design, the corrections required to use an rmANOVA model may not be able to provide a reasonable adjustment that makes the model valid.

In the case of LMEMs, one key advantage over rm-ANOVA is that they allow different structures for the variance-covariance matrix including exponential, autoregressive of order 1, rational quadratic and others^15^. Nevertheless, the analysis required to determine an appropriate variance-covariance structure for the data can be a challenging process by itself. Overall, the spherical assumption for rm-ANOVA may not capture the natural variations of the correlation in the data, and can bias the inferences from the analysis.

### 3.4 Unbalanced data

In a longitudinal study, it is frequently the case that the number of observations is different across subjects. In biomedical research, this imbalance in sample size can be caused by reasons beyond the control of the investigator (such as dropout from patients in clinical studies and attrition or injury of animals in preclinical research) leading to what is known as “missing”, “incomplete”, or (more generally speaking) unbalanced data^45^. The rm-ANOVA model is very restrictive in these situations as it assumes that observations exist for all subjects at every time point; if that is not the case subjects with one or more missing observations are excluded from the analysis. This is inconvenient because the remaining subjects might not accurately represent the population and statistical power is affected by this reduction in sample size^46^.

On the other hand, LMEMs and GAMs can work with missing observations, and inferences from the model are valid when the imbalance in the observations are *missing at random* (MAR) or *completely missing at random* (MCAR)^40,42^. In a MAR scenario, the pattern of the missing information is related to some variable in the data, but it is not related to the variable of interest^47^. If the data are MCAR, this means that the missingness is completely unrelated to the collected information^48^. Missing observations can also be *missing not at random* (MNAR) and in the case the missing observations are dependent on their value. For example, if attrition occurs in all mice that had lower weights at the beginning of a chemotherapy response study, the missing data can be considered MAR because the missigness is unrelated to other variables of interest.

However, it is worth reminding that “all models are wrong”^49^ and that the ability of LMEMs and GAMs to work with unbalanced data does not make them immune to problems that can arise due to high rates of incomplete data, such as sampling bias or a drastic reduction in statistical power. Researchers must ensure that the study design is statistically sound and that measures exist to minimize missing observation rates.

### 3.5 What does the fit of an rm-ANOVA and LMEM look like? A visual representation using simulated data

To visually demonstrate the limitations of rm-ANOVA and LMEMs for longitudinal data with non-linear trends, this section presents a simulation experiment of a normally distributed response of two groups of 10 subjects each. An rm-ANOVA model (Equation (1)), and a LMEM (Equation (4)) are fitted to each group using R^38^ and the package *nlme*^50^.

Briefly, two cases for the mean response for each group are considered: in the first case, the mean response in each group is a linear function over time with different intercepts and slopes; a negative slope is used for Group 1 and a positive slope is used for Group 2 (Figure 1A). In the second case, a second-degree polynomial (quadratic) function is used for the mean response per group: the quadratic function is concave down for Group 1 and it is concave up for Group 2 (Figure 1D). In both the linear and quadratic simulated data, the groups start with the same mean value in order to simulate the expected temporal evolution of some physiological quantity, starting at a common initial condition.

**Figure 1:**
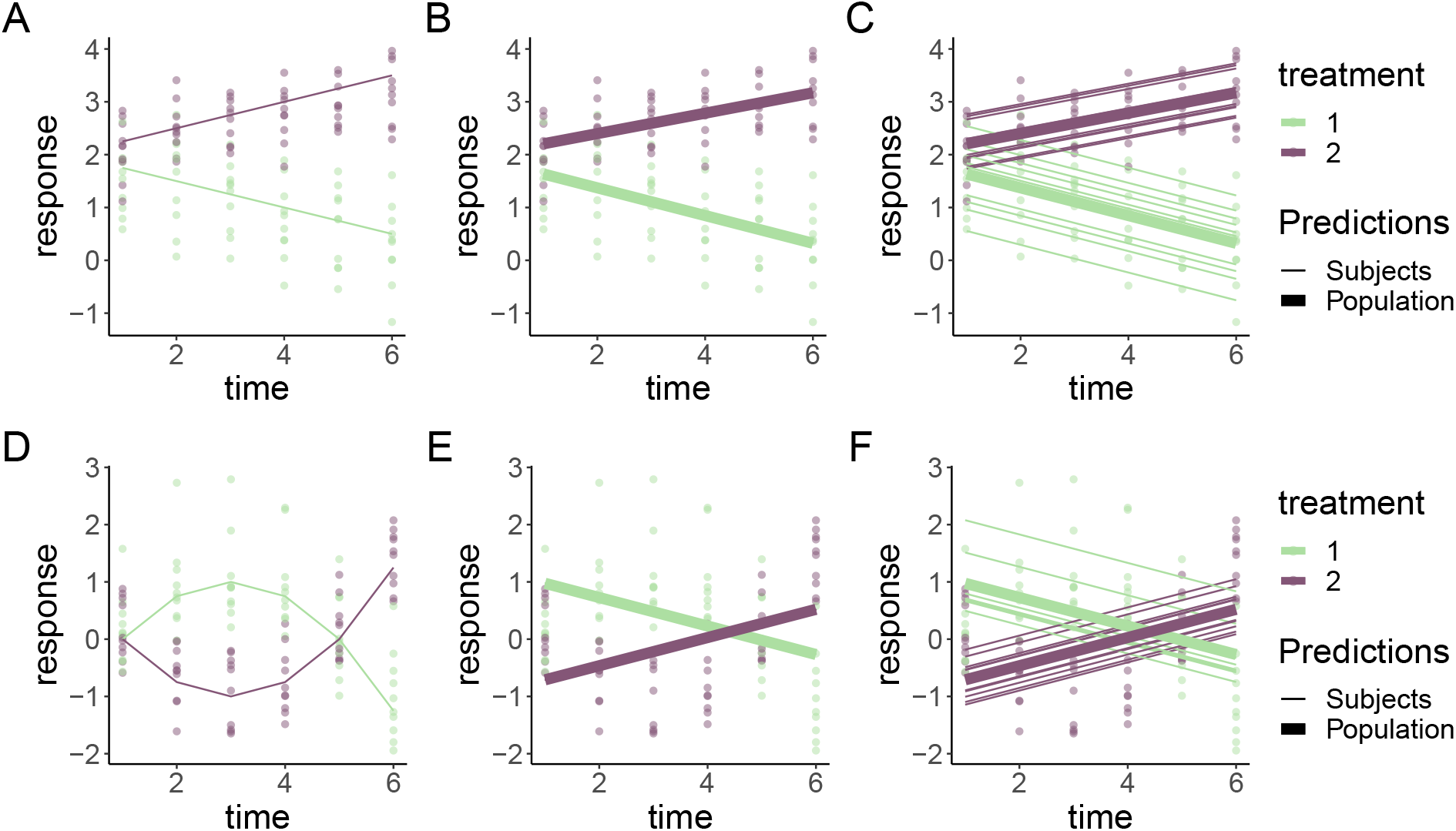
Simulated responses from two groups with correlated errors using a LMEM and a rm-ANOVA model. Top row: linear response, bottom row: quadratic response. **A**: Simulated linear data with known mean response (thick lines) and individual responses (points) showing the dispersion of the data. **D**: Simulated quadratic data with known mean response (thick lines) and individual responses (points) showing the dispersion of the data. **B**,**E**: Estimates from the rm-ANOVA model for the mean group response (linear of quadratic). Points represent the original raw data. The rm-ANOVA model not only fails to pick the trend of the quadratic data (E) but also assigns a global estimate that does not take into account the between-subject variation. **C, F**: Estimates from the LMEM in the linear and quadratic case (subject: thin lines, population: thick lines). The LMEM incorporates a random effect for each subject, but this model and the rm-ANOVA model are unable to follow the trend of the data and grossly bias the initial estimates for each group in the quadratic case (bottom row).

Specifically, the rationale for the chosen linear and quadratic functions is the expectation that a measured response in two treatment groups is similar in the initial phase of the study, but as therapy progresses a divergence in the trend of the response indicates a treatment effect. In other words, Group 1 can be thought as a “Control” group and Group 2 as a “Treatment” group. From the mean response per group (linear or quadratic), the variability or “error” of individual responses within each group is simulated using a covariance matrix with compound symmetry (constant variance across time). Thus, the response per subject at each timepoint in both the linear and quadratic simulation corresponds to the mean response per group plus the error (represented by the points in Figure 1 A, D).

A more comprehensive exploration of the fit of rm-ANOVA and LMEMs for linear and non-linear longitudinal data can be obtained from the code that appears in Appendix B, (Figures B.1 and B.2), where a simulation with compound symmetry and independent errors (errors generated from a normal distribution that are not constant over time) is presented. We are aware that the simulated data used in this section present an extreme case that might not occur frequently in biomedical research, but they are used to 1) present the consequences of modeling non-linear trends in data with a linear model such as rm-ANOVA or a LMEM with “default” (linear) effects and, 2) demonstrate that a visual assessment of model fit is an important tool that helps determine the validity of any statistical assumptions. In Section 5 we use simulated data that does follow reported trends in the biomedical literature to implement GAMs.

The simulation shows that the fits produced by the LMEM and the rm-ANOVA model are good for linear data (1B), as the predictions for the mean response are reasonably close to the “truth” of the simulated data (Figure 1A). Note that because the LMEM incorporates *random effects*, is able to provide estimates for each subject and a “population” estimate (Figure 1C).

However, consider the case when the data follows a non-linear trend, such as the simulated data in Figure 1D. Here, the mean response per group was simulated using a quadratic function, and errors and individual responses were produced as in Figure 1A. The mean response in the simulated data with quadratic behavior changes in each group through the timeline, and the mean value is the same as the initial value by the fifth time point for each group. Fitting an rm-ANOVA model (Equation (1)) or a LMEM (Equation (4)) to this data produces the fit that appears in Figure 1E, F.

Comparing the fitted responses of the LMEM and the rm-ANOVA models used in the simulated quadratic data (Figure 1E, F) indicates that the models are not capturing the changes within each group. Specifically, note that the fitted mean response of both models shows that the change (increase for Treatment 1 or decrease for Treatment 2) in the response through time points 2 and 4 is not being captured.

The LMEM is only able to account for between-subject variation by providing estimates for each subject (Figure 1F), but both models are unable to capture the fact that the initial values are the same in each group, and instead fit non-parallel lines that have initial values that are markedly different from the “true” initial values in each case (compare Figure 1D with Figure 1E, F). If such a change has important physiological implications, both rm-ANOVA and LMEMs omit it from the fitted mean response. Thus, even though the model correctly detects a divergence between treatment groups, the exact nature of this difference is not correctly identified, limiting valuable inferences from the data. It could be argued that a LMEM with quadratic effects should have been used to fit the data in Figure1F. However, because in reality the true function is not known, choosing a polynomial degree causes more questions (e.g., is it quadratic?, cubic?, or a higher degree?). Additionally, polynomial effects have other limitations, which we cover in Section 4.2.1.

This section has used simulation to better convey and visualize the limitations of linearity and correlation in the response in data with non-linear trends using an rm-ANOVA model and a LMEM, where the main issue is the expected linear trend in the response. Notice that the model misspecification is easily noticeable if the model fit and the response are visualized. In the following section, we provide a brief overview of linear models, general linear models and generalized linear mixed models before presenting the theory of GAMs, a class of semi-parametric models that can fit non-linear trends in data and that overcome the limitations of rm-ANOVA and LMEMs in the analysis of biomedical longitudinal data.

## 4 Linear Models, and beyond

Linear models (LMs) are those that assume a normal (Gaussian) distribution of the errors, and only incorporate *fixed effects* (such as rm-ANOVA). These are by far the models most commonly used to analyze data within the biomedical research community. On the other hand, Linear Mixed Effect Models (LMEMs) also incorporate *random effects*, as it has been described in Section 3.2.2.

In reality, rm-ANOVA and LMEMs are just *special cases* of a broader class of models (General Linear Models and Generalized Linear Mixed Models, respectively). In order to fully capture the constraints of such models and to understand how GAMs overcome those limitations this section will briefly provide an overview of the different classes of models and indicate how rm-ANOVA, LMEMs, and GAMs fit within this framework.

### 4.1 Generalized Linear Models (GLMs)

A major limitation of LMs is their assumption of normality in the errors. If the residuals are non-normal, a transformation is necessary in order to properly fit the model. However, transformation can lead to poor model performance^51^, and can cause problems with the biological interpretation of the model estimates. McCullagh and Nelder^52^ introduced General Linear Models (GLMs) as an extension of LMs, where the errors do not need to be normally distributed. To achieve this, consider the following model

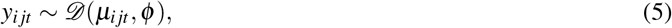

where *y*_*i jt*_ is the observation *i* in group *j* at time *t*, that is assumed to come from some distribution of the exponential family *𝒟*, with some mean *μ*_*i jt*_, and potentially, a dispersion parameter *ϕ* (which in the Gaussian case is the variance *σ*^2^). The mean (*μ*_*i jt*_) is also known as the *expected value* (or *expectation*) *E*(*y*_*i jt*_) of the observed response *y*_*i jt*_.

Then, the *linear predictor η*, which defines the relationship between the mean and the covariates can be defined as

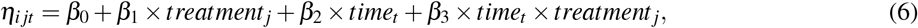

where *η*_*i jt*_ is the linear predictor for each observation *i* in each group *j*, at each timepoint *t*. Following the notation from Equation (1) the model parameters for each group are *β*_0_ (the intercept), *β*_1_, *β*_2_, and *β*_3_; *time*_*t*_ represents the covariates from each subject in each group at each time point, *treatment*_*j*_ represents the different treatment levels, and *time*_*t*_ *×treatment*_*j*_ represents their interaction.

Finally,

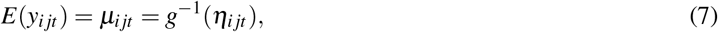

where *E*(*y*_*i jt*_) is the expectation, and *g*^−1^ is the inverse of a *link function* (*g*). The link function transforms the values from the response scale to the scale of the linear predictor *η* (Equation (6)). Therefore, it can be seen that LMs (such as rm-ANOVA) are a special case of GLMs where the response is normally distributed.

### 4.2 Generalized linear mixed models (GLMMs)

Although GLMs relax the normality assumption, they only accommodate fixed effects. Generalized Linear Mixed Models (GLMMs) are an extension of GLMs that incorporate *random effects*, which have an associated probability distribution^53^. Therefore, in GLMMs the linear predictor takes the form

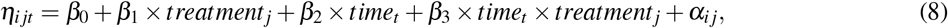

where *α*_*i j*_ corresponds to the random effects that can be estimated within each subject in each group, and all the other symbols correspond to the notation of Equation (6). Therefore, LMEMs are special case of GLMMs where the distribution of the response is normally distributed^52^, and GLMs are a special case of GLMMs where there are no random effects. In-depth and excellent discussions about LMs, GLMs and GLMMs can be found in Dobson^54^ and Stroup^55^.

#### 4.2.1 GAMs as a special case of Generalized Linear Models

##### 4.2.1.1 GAMs and Basis Functions

Notice that in the previous sections, the difference between GLMs and GLMMs resides on their linear predictors (Equations (6), (8)). Generalized additive models (GAMs) are an extension of the GLM family that allow the estimation of smoothly varying trends where the relationship between the covariates and the response is modeled using *smooth functions*^34,37,56^. In a GAM, the linear predictor has the form

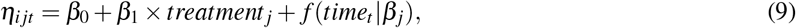

where *β*_0_ is the intercept, and *β*_1_ is the coefficient for each treatment group. Notice that the construction of the predictor is similar to that of Equation (6), but in this case the parametric terms involving the effect of time, and the interaction between time and treatment have been replaced by the smooth term *f* (*time*_*t*_|*β* _*j*_). The smooth term *f* (*time*_*t*_|*β* _*j*_) gives a different smooth response for each treatment. ^1^ A GAM version of a linear model can be written as

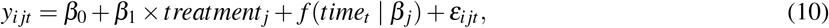

where *y*_*i jt*_ is the response at time *t* of subject *i* in group *j*, and *ε*_*i jt*_ represents the deviation of each observation from the mean.

In contrast to the linear functions used to model the relationship between the covariates and the response in rmANOVA or LMEM, the use of smooth functions in GAMs is advantageous as it does not restrict the model to a linear relationship, although a GAM can estimate a linear relationship if the data is consistent with a linear response. One possible set of functions for *f* (*time*_*t*_ | *β*_*j*_) that allow for non-linear responses are polynomials (which can also be used in LMEMs), but a major limitation is that polynomials create a “global” fit as they assume that the same relationship exists everywhere, which can cause problems with inference^36^. In particular, polynomial fits are known to show boundary effects because as *t* goes to *±*∞, *f* (*time*_*t*_ | *β*_*j*_) goes to *±*∞ which is almost always unrealistic and causes bias at the endpoints of the time period.

The smooth functional relationship between the covariates and the response in GAMs is specified using a semiparametric relationship that can be fit within the GLM framework, using a *basis function* expansion of the covariates and estimating random coefficients associated with these basis functions. A *basis* is a set of functions that spans the mathematical space within which the true but unknown *f* (*time*_*t*_) is thought to exist^34^. For the linear model in Equation (1), the basis coefficients are *β*_1_, *β*_2_ and *β*_3_ and the basis vectors are *treatment*_*j*_, *time*_*t*_, and *time*_*t*_ *×treatment*_*j*_. The basis function then, is the linear combination of basis coefficients and basis vectors that map the possible relationship between the covariates and the response^57^, which in the case of Equation (1) is restricted to a linear family of functions. In the case of Equation (10), the basis functions are contained in the expression *f* (*time*_*t*_ | *β*_*j*_), which means that the model allows for non-linear relationships among the covariates.

Splines (which derive their name from the physical devices used by draughtsmen to draw smooth curves) are commonly used as basis functions that have a long history in solving semi-parametric statistical problems and are often a default choice to fit GAMs as they are a simple, flexible, and powerful option to obtain smoothness^58^. Although different types of splines exist, cubic, thin plate splines, and thin plate regression splines will be briefly discussed next to give a general idea of these type of basis functions, and their use within the GAM framework.

Cubic splines (CS) are smooth curves constructed from cubic polynomials joined together in a manner that enforces smoothness. The use of CS as smoothers in GAMs was discussed within the original GAM framework^56^, but they are limited by the fact that their implementation requires the selection of some points along the covariates (known as ‘knots’, the points where the basis functions meet) to obtain the finite basis, which affects model fit^59^. A solution to the knot placement limitation of CS is provided by thin plate splines (TPS), which provide optimal smooth estimation without knot placement, but that are computationally costly to calculate^37,59^. In contrast, thin plate regression splines (TPRS) provide a reasonable “low rank” (truncated) approximation to the optimal TPS estimation, which can be implemented in an computationally efficient^59^. Like TPS, TPRS only requires specifying the number of basis functions to be used to create the smoother (for mathematical details on both TPS and TPRS see Wood^37,9^).

To further clarify the concept of basis functions and smooth functions, consider the simulated response for Group 1 that appears in Figure 1D. The simplest GAM model that can be used to estimate such response is that of a single smooth term for the time effect; i.e., a model that fits a smooth to the trend of the group through time. A computational requisite in *mgcv* is that the number of basis functions to be used to create the smooth cannot be larger than the number of unique values from the independent variable. Because the data has six unique time points, we can specify a maximum of six basis functions (including the intercept) to create the smooth. It is important to note that is not necessary to specify a number of basis equal to the number of unique values in the independent variable; fewer basis functions can be specified to create the smooth as well, as long as they reasonably capture the trend of the data.

Here, the main idea is that the resulting smooth matches the data and approximates the true function without becoming too “wiggly” due to the noise present. A detailed exploration of wiggliness and smooth functions is beyond the scope of this manuscript, but in essence controlling the wiggliness (or “roughness”) of the fit is achieved by using a *smoothness parameter* (*λ*), which is used to penalize the likelihood by multiplying it with the integrated square of the second derivative of the spline smooth. The second derivative of the spline smooth is a measure of curvature, or the rate of change of the slope^34,37^, and increasing the penalty by increasing *λ* results in models with less curvature. As *λ* increases, the parameter estimates are penalized (shrunk towards 0) where the penalty reduces the wiggliness of the smooth fit to prevent overfitting. In other words, a low penalty estimate will result in wiggly functions whereas a high penalty estimate provides evidence that a linear response is appropriate.

With this in mind, if four basis functions (plus the intercept) are used to fit a GAM for the data of Group 1 (concave down trend) that appears in Figure 1D, the resulting fitting process is shown in Figure 2. In Figure 2A the four basis functions (and the intercept) are shown. Each of the five basis functions is evaluated at six different points (because there are six points on the timeline). The coefficients for each of the basis functions of Figure 2A are estimated using a penalized regression with smoothness parameter *λ*, that is estimated when fitting the model. The penalized coefficient estimates fitted are shown in Figure 2B.

**Figure 2:**
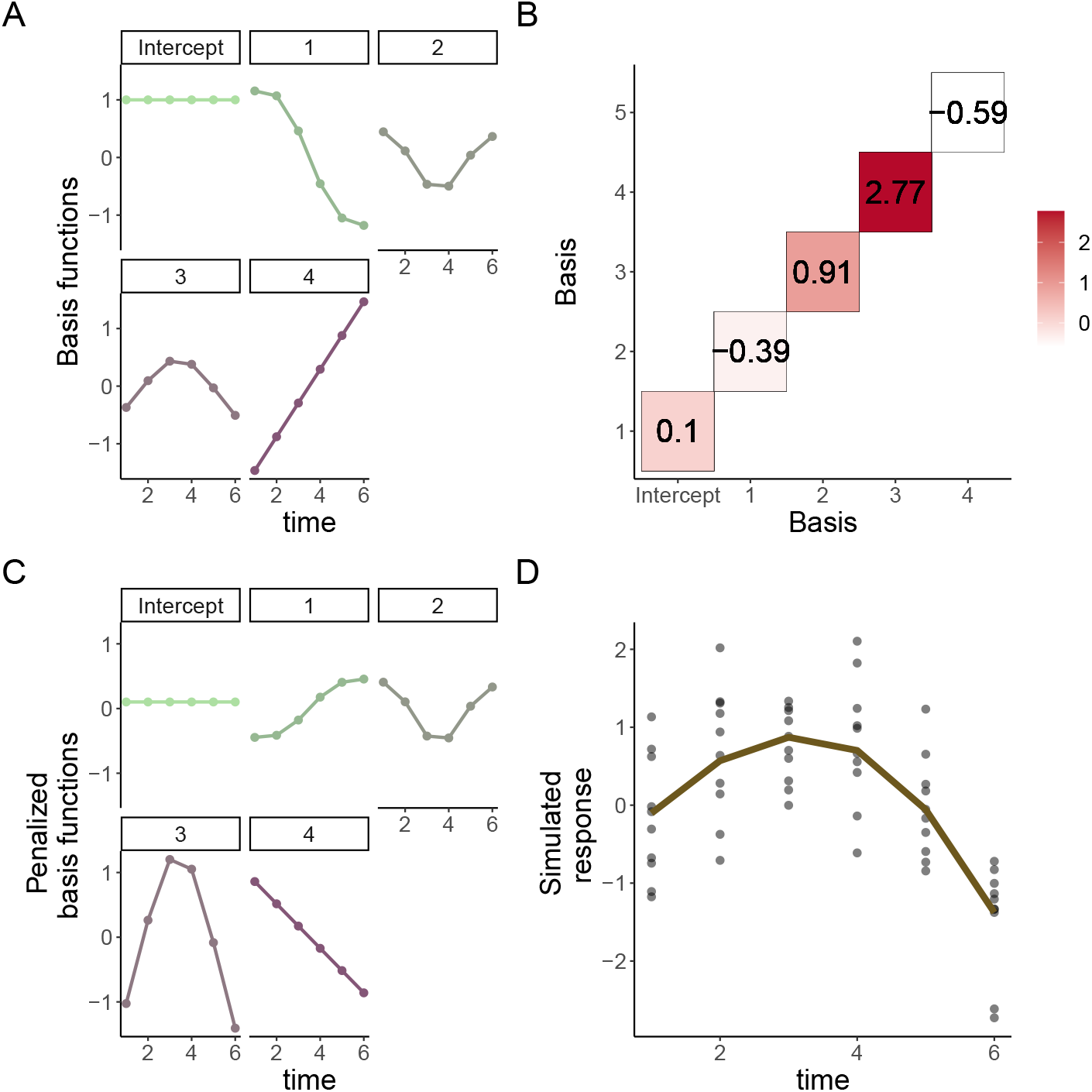
Basis functions for a single smoother for time. **A**: Basis functions for a single smoother for time for the simulated data of Group 1 from Figure 2. **B**: Matrix of basis function weights. Each basis function is multiplied by a coefficient which can be positive or negative. The coefficient determines the overall effect of each basis in the final smoother. **C**: Weighted basis functions. Each of the four basis functions (and the intercept) of panel A has been weighted by the corresponding coefficient shown in Panel B. Note the corresponding increase (or decrease) in magnitude of each weighted basis function. **D**: Smoother for time and original data points. The smoother (line) is the result of the sum of each weighted basis function at each time point, with simulated values for Group 1 shown as points.

To get the weighted basis functions, each basis (from Figure 2A) is multiplied by the corresponding coefficients in Figure 2B, thereby increasing or decreasing the original basis functions. Figure 2C shows the resulting weighted basis functions. Note that the magnitude of the weighting for the first basis function has resulted in a decrease of its overall contribution to the smoother term (because the coefficient for that basis function is negative and its magnitude is less than one). On the other hand, the third basis function has roughly doubled its contribution to the smooth term. Finally, the weighted basis functions are added at each timepoint to produce the smooth term. The resulting smooth term for the effect of *time* is shown in Figure 2D (brown line), along the simulated values which appear as points.

#### 4.2.2 A Bayesian interpretation of GAMs

Bayes’ theorem states that the probability of an event can be calculated using prior knowledge and observed data^60^. In the case of data that shows non-linear trends, the prior that the *true* trend of the data is likely to be smooth rather than “wiggly” introduces the concept of a prior distribution for wiggliness (and therefore a Bayesian view) of GAMs^37^. Moreover, GAMs are considered “empirical” Bayesian models when fitted using the package *mgcv* because the smoothing parameters are estimated from the data (and not from a posterior distribution as in the “fully Bayesian” case, which can be fitted using JAGS, Stan, or other probabilistic programming language)^61^. Therefore, the confidence intervals (CIs) calculated by default for the smooth terms using *mgcv* are considered empirical Bayesian credible intervals^33^, which have good *across the function* (“frequentist”) coverage^37^.

To understand across the function coverage, recall that a CI provides an estimate of the region where the “true” or “mean” value of a function exists, taking into account the randomness introduced by the sampling process. Because random samples from the population are used to calculate the “true” value of the function, there is inherent variability in the estimation process and the CI provides a region with a nominal value (usually, 95%) where the function is expected to lie. In an *across the function* CI (like those estimated by default for GAMs using *mgcv*), if we average the coverage of the interval over the entire function we get approximately the nominal coverage (95%). In other words, we expect that about 95% of the points that compose the true function will be covered by the across the function CI. As a consequence, some areas of the CI for the function have more than nominal coverage and some areas less than the nominal coverage.

Besides the across the function CI, “simultaneous” or “whole function” CIs can also be computed, which contain the *whole function* with a specified probability^37^. Suppose we chose a nominal value (say, 95%) and compute a simultaneous CI; if we obtain 100 repeated samples and compute a simultaneous CI in each case, we would expect that the true function lies completely within the computed simultaneous CI in 95 of those repeated samples. Briefly, to obtain a simultaneous CI we simulate 10,000 draws from the empirical Bayesian posterior distribution of the fitted smooths. Then, we obtain the maximum absolute standardized deviation of the differences in smooth estimates which is used to correct the coverage of the across the function CI^62^ in a similar fashion to how *q-values* correct *p-values* to control false positive discovery rates^63^.

In-depth theory of the Bayesian interpretation of GAMs and details on the computation of simultaneous and across the function CIs are beyond the scope of this paper, but can be found in Miller^61^, Wood^37,^ Simpson^34^, Marra^64^, and Ruppert^62^. What we want to convey is that a Bayesian interpretation of GAMs allows for robust estimation using simultaneous empirical Bayesian CIs, as their estimates can be used to make comparisons between different groups in a similar way that multiple comparisons adjustments make inference from ANOVA models more reliable.

With this in mind, in the next section we consider the use of GAMs to analyze longitudinal biomedical data with non-linear trends, and use simultaneous empirical Bayesian CIs to assess significance between treatment groups.

## 5 The analyisis of longitudinal biomedical data using GAMs

The previous sections provided the basic understanding of the GAM framework and how these models are more advantageous to analyze non-linear longitudinal data when compared to rm-ANOVA or LMEMs. This section will use simulation to present the practical implementation of GAMs for longitudinal biomedical data using R and the package `mgcv`. A brief guide for model selection and diagnostics appears in Appendix A, and the code for the simulated data and figures can be found in Appendix B.

### 5.1 Simulated data

The simulated data is based on the reported longitudinal changes in oxygen saturation (StO_2_) in subcutaneous tumors (Figure 3C in Vishwanath et. al.^16^), where diffuse reflectance spectroscopy was used to quantify StO_2_ changes in both groups at the same time points (days 0, 2, 5, 7 and 10). In the “Treatment” group (chemotherapy) an increase in StO_2_ is observed through time, while a decrease is seen in the “Control” (saline) group. Following the reported trend, we simulated 10 normally distributed observations at each time point with a standard deviation (SD) of 10% (matching the SD in the original paper). The simulation based on the real data appears in Figure 3A.

**Figure 3:**
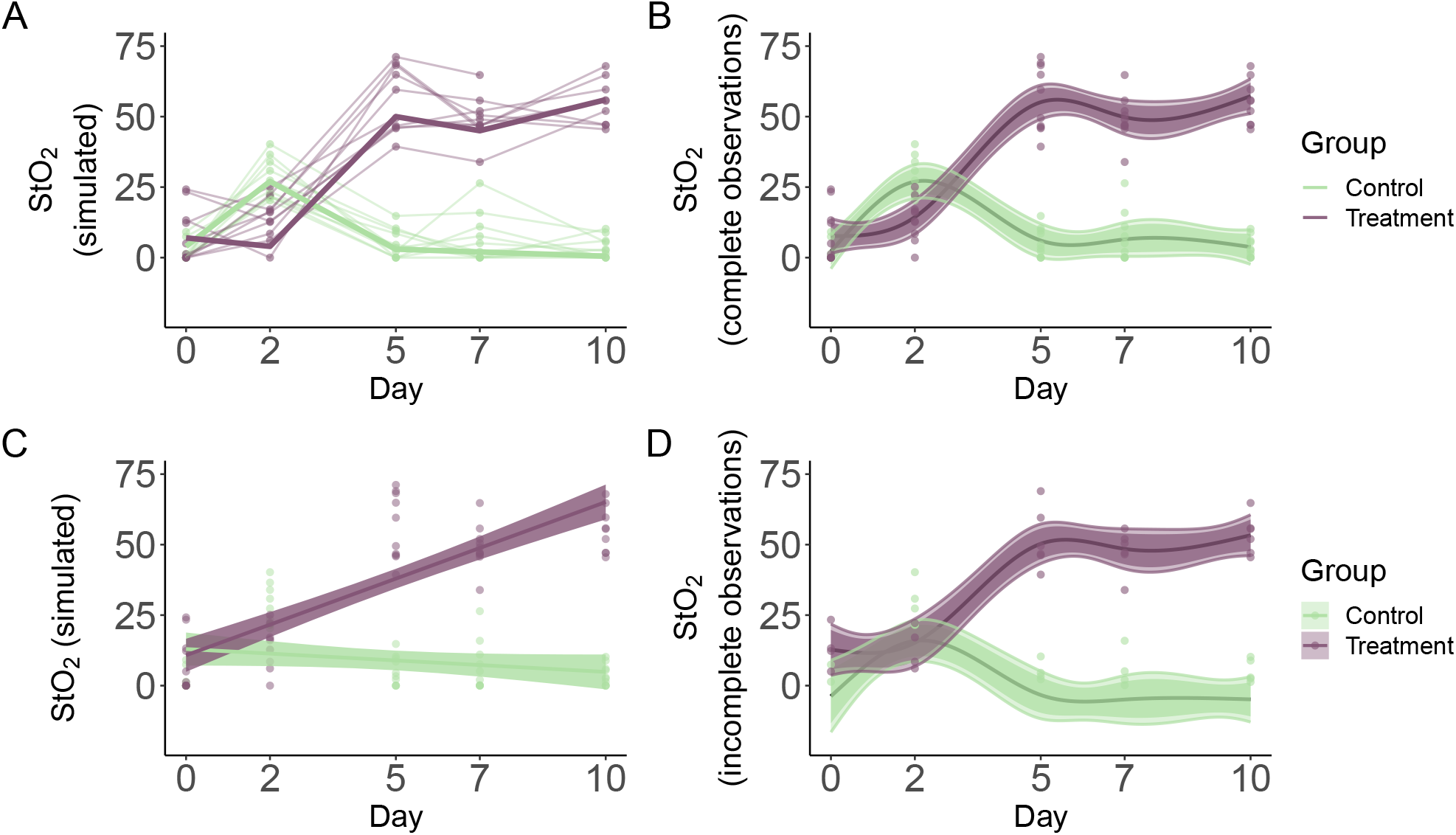
Simulated data and smooths for oxygen saturation in tumors. **A**: Simulated data (thin lines) that follows previously reported trends (thick lines) in tumors under chemotherapy (Treatment) or saline (Control) treatment. Simulated data is from a normal distribution with standard deviation of 10% with 10 observations per time point. **B**: Smooths from the GAM model for the full simulated data with interaction of Group and Treatment. Lines represent trends for each group, shaded regions are 95% across the function (narrow region) and simultaneous (wide region) confidence intervals. **C**: The rm-ANOVA model for the simulated data, which does not capture the changes in each group over time. **D**: Smooths for the GAM model for the simulated data with missing observations (40%). Lines represent trends for each group, shaded regions are 95% across the function (narrow region) and simultaneous (wide region) confidence intervals.

### 5.2 An interaction GAM for longitudinal data

An interaction effect is typically the main interest in longitudinal biomedical data, as the interaction takes into account treatment, time, and their combination. In a practical sense, when a GAM is implemented for longitudinal data, a smooth can be added to the model for the *time* effect for each treatment to account for the repeated measures over time. Although specific methods of how GAMs model correlation structures is a topic beyond the scope of this paper, it suffices to say that GAMs are flexible and can handle correlation structures beyond compound symmetry. A detailed description on the close relationship between basis functions and correlation functions can be found in Hefley et. al.^57^.

For the data in Figure 3A, the main effect of interest is how StO_2_ changes over time for each treatment. To estimate this, the model incorporates separate smooths for each *Group* as a function of *Day*. The main thing to consider is that model syntax accounts for the fact that one of the variables is numeric (*Day*) and the other is a factor (*Group*). Because the smooths are centered at 0, the factor variable needs to be specified as a parametric term in order to identify any differences between the group means. Using R and the package `mgcv` the model syntax is:

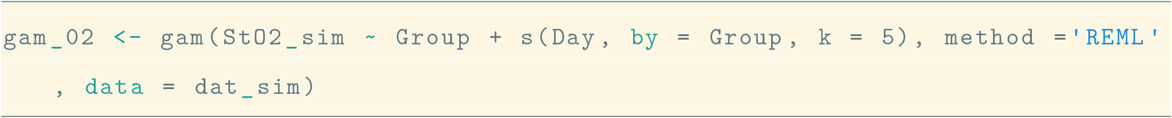

This syntax specifies that `gam_02` (named this way so it matches the model workflow from Appendix A) contains the fitted model, and that the change in the simulated oxygen saturation (`StO2_sim`) is modeled using independent smooths over `Day` for each `Group` (the parenthesis preceded by `s`) using four basis functions (plus intercept). The smooth is constructed by default using TPRS, but other splines can be used if desired, including Gaussian process smooths^34^ (a description of all the available smooths can be found by typing `?mgcv::smooth.terms` in the Console). Finally, the parametric term `Group` is added to quantify overall mean differences in the effect of treatment between groups, as we have indicated above.

Although the default `method` used to estimate the smoothing parameters in *mgcv* is generalized cross validation (GCV), Wood^37^ showed the restricted maximum likelihood (REML) to be more resistant to overfitting while also easing the quantification of uncertainty in the smooth parameters; therefore in this manuscript REML is always used for smooth parameter estimation. An additional argument (`family`) allows to specify the expected distribution of the response, but it is not used in this model because we expect a normally-distributed response (which is the default `family` in *mgcv*).

When the smooths are plotted over the raw data, it is clear that the model has been able to capture the trend of the change of StO_2_ for each group across time (Figure 3B). Model diagnostics can be obtained using the `gam.check` function, and the function `appraise` from the package *gratia*^65^ as we show in Appendix A. Additional discussions on model selection can be found in Wood^37^ and Harezlak^66^.

One question that might arise at this point is “what is the fit that an rm-ANOVA model produces for the simulated data?”. The fit of an rm-ANOVA model, which corresponds to Equation (1), is presented in Figure 3C. This is a typical case of model misspecification: The slopes of each group are different, which would lead to a *p-value* indicating significance for the treatment and time effects, but the model is not capturing the changes that occur at days 2 and between days 5 and 7, whereas the GAM model is able to reliably estimate the trend over all timepoints (Figure 3B).

Because GAMs do not require equally-spaced or complete observations for all subjects (as rm-ANOVA does), they are advantageous to analyze longitudinal data where unbalanced data exists. The rationale behind this is that GAMs are able to pick the trend in the data even when some observations are incomplete. However, this usually causes the resulting smooths to have wider confidence intervals and less ability to discern differences in trends. To exemplify this, consider the random deletion of 40% of the simulated StO_2_ values from Figure 3A. If the same interaction GAM (`gam_02`) is fitted to this data with unbalanced observations, the resulting smooths appear in (Figure 3D). Note that the model is still able to show a different trend for each group, but with a somewhat more linear profile in some areas.

Additionally, note that in Figure 3B,D we show two CIs for each of the fitted smooths (shaded regions). The across the function CIs are represented by the narrow regions and because the simultaneous CIs contain the whole function on a nominal value, they are wider than the across the function CI, resulting in the wide shaded regions. For the dataset with incomplete observations, the CIs for the smooths overlap during the first 3 days because the estimates are less robust with fewer data points, and the trend is less pronounced than in the full dataset. However, the overall trend of the data is picked by the model in both cases, with as few as 4 observations per group at certain time points.

### 5.3 Determination of significance in GAMs for longitudinal data

At the core of a biomedical longitudinal study lies the question of a significant difference between the effect of two or more treatments in different groups. In linear models (such as rm-ANOVA), if there is a significant *p-value* after a *post-hoc* analysis we then can make inference about the effect size using the slope or the intercept from the model. In GAMs however, there is no single *p-value* to determine the significance of an effect as in linear models. Therefore, the coefficients of GAMs do not provide a simple interpretation as in the linear model case, but the changes in slope at specific timepoints can be used to determine the instantaneous effect size. In essence, the idea behind the estimation of significance in GAMs across different treatment groups is that the difference between the separate smoothers per group (such as in `gam_02`) can be computed pairwise, followed by the estimation of an empirical Bayesian simultaneous CI around this difference.

The pairwise difference in smooths can be conceptualized in the following manner: Different time trends in each group are an indication of an effect by the treatment as in Figure 3A, where the chemotherapy causes StO_2_ to increase over time. With this expectation of different trends in each group, computing the difference between the trends will identify if the observed difference is significant. A difference between groups with similar trends is unlikely to be distinguishable from zero, which would indicate that the treatment is not causing a change in the response in one of the groups (assuming the other group is a Control or Reference group). Therefore, the computation of both the difference between the smooths and the corresponding simultaneous empirical Bayesian CI around this difference is able to provide an estimation of when and by how much there is a significant difference between the different groups. Additionally, the correction provided by the simultaneous empirical Bayesian CI makes the estimation robust as we know that on average, the simultaneous CI will contain the *whole function* at a nominal value (say, 95%).

To visualize this, consider the calculation of pairwise differences for the fitted smooths that appear in Figure 3B, D. Figure 4 shows the difference between each treatment group trend for the full and missing datasets with a simultaneous CI computed around the difference. Here, the “Control” group is used as the reference to which “Treatment” group is being compared. Notice that because we have included the means of each group, there is correspondence between the scale of the original data and the scale of the pairwise comparisons. This can be seen in Figure 3B, where at day 5 there is essentially a difference of 50% between StO_2_ in both groups, which corresponds to the -50% difference in 4A. However, if there are multiple parametric terms in the model (more factors that need to be specified), such inclusion of the means can become problematic. However, we believe that the model we have presented here suffices in a wide range of situations where adding the group means is a relatively easy implementation that can help better visualize the estimation from the model from a biological perspective.

**Figure 4:**
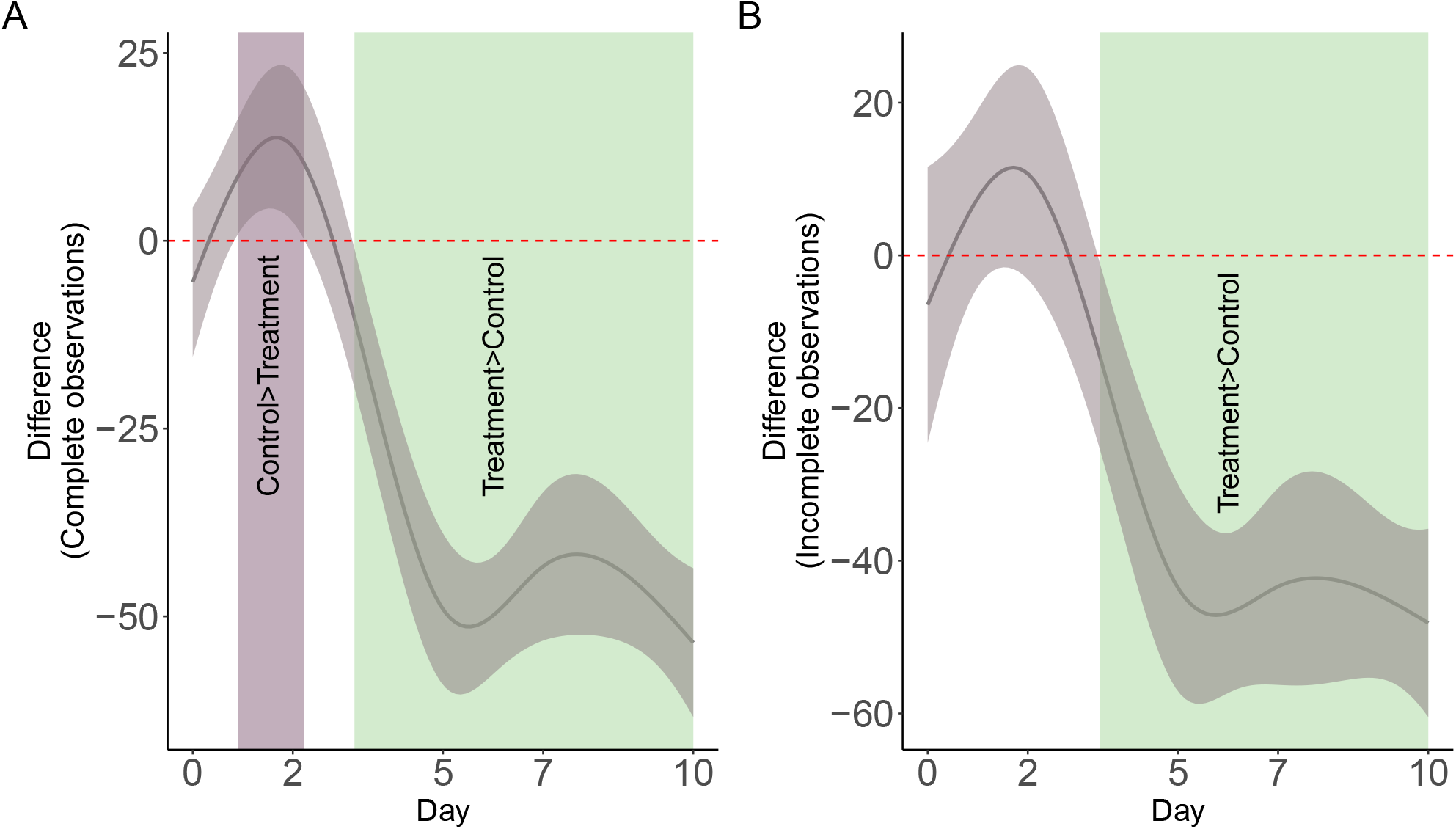
Pairwise comparisons for smooth terms. **A**: Pairwise comparisons for the full dataset. **B**: Pairwise comparisons for the dataset with incomplete observations. Significant differences exist where the 95% empirical Bayesian simultaneous CI does not cover 0. In both cases the effect of treatment is significant after day 3. For the difference, we have included the means so the value of the difference has direct correspondence with the scale of the response (Figure 3A).

In Figure 4A, the shaded regions over the confidence interval (where the CI does not cover 0) indicate the time interval where each group has a different mean effect than the other. Notice that the shaded region between days 1 and ≈ 2 for the full dataset indicates that through that time, the “Control” group has higher mean StO_2_, but as therapy progresses the effect is reversed and by day ≈ 3 it is the “Treatment” group that statistically on average has greater StO_2_. This would suggest that the effect of chemotherapy in the “Treatment” group becomes significant after day 3 for the given model. Moreover, notice that although there is no actual measurement at day 3, the model is capable of providing an estimate of when the shift in mean StO_2_ occurs.

On the data with missing observations (Figure 3D), the smooth pairwise comparison (Figure 4B) shows that because the confidence intervals overlap zero for the first two days there is no statistically significant difference between the groups. However, because the model is still able to pick the overall trend in StO_2_, the pairwise comparison is able to estimate the change on day 3 where the mean difference between groups becomes statistically significant as in the full dataset smooth pairwise comparison.

For biomedical studies, the ability of smooth comparisons to provide an estimate of *when* and by *how much* a biological process becomes significant is advantageous because it can help researchers gain insight on metabolic changes and other biological processes that can be worth examining, and can help refine the experimental design of future studies in order to obtain measurements at time points where a significant change might be expected.

## 6 Discussion

Biomedical longitudinal data is particularly challenging to analyze due to the frequency of incomplete observations and different correlation structures in the data, which limit the use of rm-ANOVA. Although LMEMs can handle unbalanced observations and different correlation structures, both LMEMs and rm-ANOVA yield biased estimates when they are used to fit data with non-linear trends as we have visually demonstrated in Section 3.5. When these models do not capture the non-linear trend of the data, this results in a “model misspecification error”. This “model misspecification” error, also is known as a “Type III” error^17^ is particularly important because although the *p-value* is the common measure of statistical significance, the validity of its interpretation is determined by the agreement of the data and the model.

Guidelines for statistical reporting in biomedical journals exist (the SAMPL guidelines)^67^ but they have not been widely adopted and in the case of longitudinal data, we consider that researchers would benefit from reporting a visual assessment of the correspondence between the model fit and the data, rather than merely relying on a *R*^2^ or *p-value* value, whose interpretation is not clear in the case of a Type III error.

In this paper we have presented GAMs as a suitable method to analyze longitudinal data with non-linear trends. It is interesting to note that although GAMs are a well established method to analyze temporal data in different fields (e.g., which are palaeoecology, geochemistry, and ecology)^33,57^ they are not routinely used in biomedical research despite an early publication from Hastie and Tibshirani that demonstrated their use in medical research^68^. This is possibly due to the fact that the theory behind GAMs can seem very different from that of rm-ANOVA and LMEMs.

However, in Section 4.2.1 we demonstrated that at its core the principle underlying GAMs is quite simple: Instead of using a linear relationship to model the response (as rm-ANOVA and LMEMs do), GAMs use basis functions to build smooths that are capable of learning non-linear trends in the data. The use of basis functions is a major advantage over models where the user has to know the non-linear relationship *a priori*, such as in the case of polynomial effects in LMEMs where in addition, there is no biological interpretation for such polynomial assumptions. This does not mean, however, that as any other statistical model GAMs do not have certain limitations. In particular, beyond the range of the data GAMs only reflect the assumptions built into the basis functions, be that flat values or linear extrapolation of the slope. Therefore, researchers need to be careful when using GAMs for extrapolating purposes. In addition, both polynomial and GAMs show higher variance in estimates near the boundary of the data, but additive models generally have less variance than polynomials. However, because GAMs let the data speak for themselves, they provide estimates that are consistent with non-linear trends and therefore can be used to obtain an accurate representation of the effect of time in a biological process.

Beyond the theory, from a practical standpoint is equally important to demonstrate how GAMs are computationally implemented. We have provided an example on how GAMs can be fitted using simulated data that follows trends reported in biomedical literature^16^ using R and the package *mgcv*^37^ in Section 5, while a basic workflow for model selection is in Appendix A.

One of the features of GAMs is that their Bayesian interpretation allows for inference about differences between groups without the need of a *p-value*, thereby providing a time-based estimate of shifts in the response that can be directly tied to biological values as the pairwise smooth comparisons in Figure 4 indicate. The GAM is therefore able to identify changes between the groups at time points where data was not directly measured and in the case of incomplete observations (≈ day 3 in Figure 4 A, B). This more nuanced inference can be used by researchers as feedback on experiment design and to further evaluate important biological changes in future studies.

We have used R as the software of choice for this paper because it provides a fully developed environment to fit GAMs, enables simulation (which is becoming increasingly used for exploratory statistical analysis and power calculations), and provides powerful and convenient methods of visualization, which are key aspects that biomedical researchers might need to consider to make their work more reproducible. In this regard, reproducibility is still an issue in biomedical research^69,70^, but it is becoming apparent that what other disciplines have experienced in this aspect is likely to impact this field sooner rather than later. Researchers need to plan on how they will make their data, code, and any other materials open and accessible as more journals and funding agencies recognize the importance and benefits of open science in biomedical research. We have made all the data and code used in this paper accessible, and we hope that this will encourage other researchers to do the same with future projects.

## 7 Conclusion

We have presented GAMs as a method to analyze longitudinal biomedical data. Future directions of this work will include simulation-based estimations of statistical power using GAMs, as well as demonstrating the prediction capabilities of these models using large datasets. By making the data and code used in this paper accessible, we hope to address the need of creating and sharing reproducible work in biomedical research.

## Supporting information

Appendix A: GAM workflow

## 8 Acknowledgements

This work was supported by the National Science Foundation Career Award (CBET 1751554, TJM) and the Arkansas Biosciences Institute.

## 9 Declaration of Conflicting Interests

The Authors declare that there is no conflict of interest.

## Supplementary Materials

Two Appendices which contain a basic workflow to implement GAMs in R and all the code used to create this manuscript are available as PDFs in the Supplementary Material. A GitHub repository containing all the code used for this paper along with detailed instructions for its use is available at https://github.com/aimundo/GAMs-biomedicalresearch.

## Footnotes

1 If the smooth term represented a linear relationship, then *f* (*time*_*t*_|*β*_*j*_) = *β*_2_ × *time*_*t*_ + *β*_3_ × *time*_*t*_ × *treatment*_*j*_; however, in general, the smooth term is a more flexible function than a linear relationship, with parameter vectors *β*_*j*_ for each treatment

## Notes

https://github.com/aimundo/GAMs-biomedical-research

### Competing Interest Statement

The authors have declared no competing interest.

### Summary of Updates

The comments from the peer review process significantly improved the scope, clarity, and organization of the manuscript. The authors believe that due to these changes, the preprint needs to be revised accordingly so it can be shared with the public at large rapidly.

