## Appendix A: GAM workflow for "Generalized additive models to analyze non-linear trends in biomedical longitudinal data using R: Beyond repeated measures ANOVA and Linear Mixed Models"

### APPENDIX A: WORKFLOW FOR GAMs FOR BIOMEDICAL LONGITUDINAL DATA

Ariel I. Mundo, John R. Tipton, and Timothy J. Muldoon*

**

This appendix shows a basic workflow to fit a series of increasingly complex GAMs to simulated data that follows the trends presented in Section 5.1 in the main manuscript. Graphical and parameter diagnostics for goodness of fit are discussed, as well as model comparison via AIC (Aikake Information Criterion). For simplicity, the confidence intervals (CIs) shown in this section for the models are the across the function CIs created by mgcv by default. However, for the pairwise comparisons of the third model we use *simultaneous intervals* as in the main manuscript, and the code for creating simultaneous CIs for the smooths can be found in Appendix B.

### A.1 Setup and data simulation

First, we load the libraries needed for all analyses and figures. We also set seed for reproducibility.

```
library(patchwork)
library(tidyverse)
library(mvnfast)
library(nlme)
library(mgcv)
library(gratia)
library(here)
library(scico)
set.seed(2021)  #set seed for reproducibility

# alpha for ribbon in the smooth plots
al <- 0.8

thm1 <- scale_fill_scico_d(palette = "tokyo", begin = 0.3, end = 0.8,
   direction = -1,
    aesthetics = c("colour", "fill"))
```

Next, we create a data frame with the same trends from the main manuscript:

- Two treatment groups (Control and Treatment)
- Five time points (days 0, 2, 5, 7,10)
- Trends in $StO_2$ for both groups

```
dat <- tibble(StO2 = c(4, 27, 3, 2, 0.5, 7, 4, 50, 45, 56), Day = rep(c(0,
    2, 5,
    7, 10), times = 2), Group = as.factor(rep(c("Control", "Treatment"),
       each = 5)))
```

We do not include a plot of the simulated data here because it follows the same trend from Figure 3A in the main manuscript (however, it can be easily plotted if desired). Finally, we call the function simulate_data in the script simulate_data.R to create 10 replicates at each time point from a normal distribution with a standard deviation of 10% in StO2.

```
n <- 10  #number of observations
sd <- 10  #approximate sd from paper
source(here("scripts", "simulate_data.R"))
dat_sim <- simulate_data(dat, n, sd)
```

### A.2 First model

The first model fitted to the data is one that only assumes a single smooth over time shared by both the treatment and control groups The model syntax specifies that gam_00 is the object that will contain all the model information, and that the model attempts to explain changes in StO2_sim (simulated $StO_2$) using a smooth per `Day`. The model will use four basis functions (`k = 5`) for the smooth. The smooth is constructed by default using thin plate regression splines. The smoothing parameter estimation method used is the restricted maximum likelihood (`REML`).

```
gam_00 <- gam(StO2_sim ~ s(Day, k = 5), method = "REML", data = dat_sim)
```

To obtain model diagnostics, two methodologies are to be used: 1) graphical diagnostics, and 2) a model check. In the first case, the functions `appraise` and `draw` from the package *gratia* can be used to obtain a single output with all the graphical diagnostics. For model check, the functions `gam.check` and `summary` from *mgcv* provide detailed information about the model fit and its parameters. Keep in mind that `gam.check` is a function that also provides the graphical diagnostics obtained using *gratia*, if such graphical output is not desired the source code can be accessed typing `gam.check` in the Console, and the code without the graphical output can be used in a custom function (which is the approach we follow later on this Appendix).

#### A.2.1 Graphical diagnostics

The following code chunk generates graphical checks for the fitted GAM model with a single shared smooth shared between the treatment and control groups.

```
appr1 <- appraise(gam_00)
sm1 <- draw(gam_00)
visual_check <- appr1 + sm1

visual_check + plot_layout(design = layout1)
```

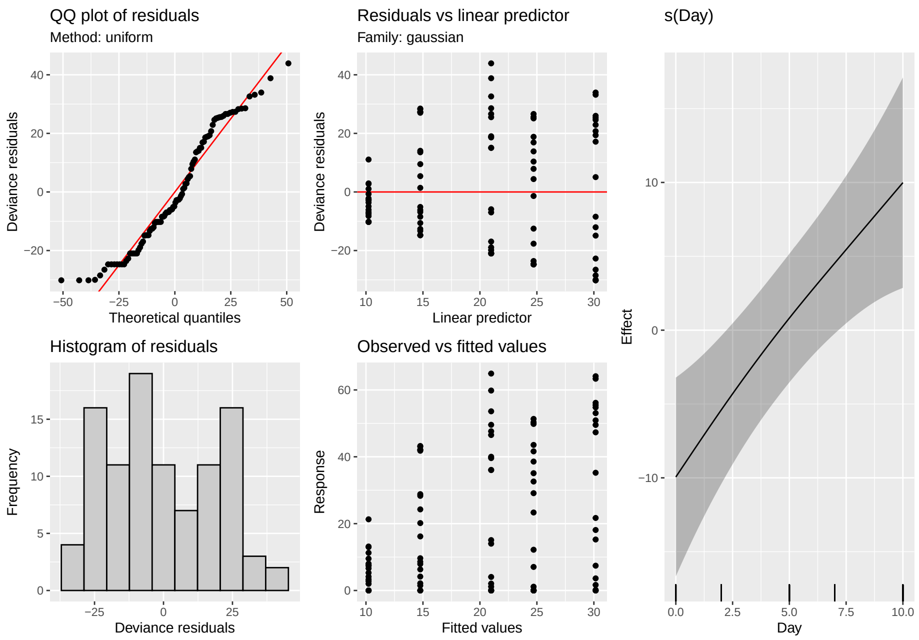


Figure A.1: Graphical diagnostics for the first GAM model. Left: Graphical diagnostics provided by the function `appraise` from the package *gratia*. Right: Fitted smooth for the model, provided by the function `draw`.

From the output of the function `appraise` in Figure A.1, the major indicators of concern about the model are the QQ plot of residuals and the histogram of residuals. The QQ plot shows that the errors are not reasonably located along the 45° line (which indicates normality), as there are multiple points that deviate from the trend, specially in the tails. The histogram also shows that the variation (residuals) is not following the assumption of a normal distribution.

The draw function permits to plot the smooths as ggplot2 objects, which eases subsequent manipulation, if desired. Because model gam_00 specifies only one smooth for the time covariate (Day), the plot only contains only one smooth. Note that the smooth shows an almost linear profile.

#### A.2.2 Model check

Special attention must be paid to the parameter 'k-index' from gam.check (which calls k.check to perform the calculation). This parameter indicates if the basis dimension of the smooth is adequate, i.e., it checks that the basis used to create the smooth are adequate to capture the trends in the data. If the model is not adequately capturing the trends in the data, this is indicated by a low k-index value (<1). Because we plot the model diagnostics using appraise later, the graphical output from gam.check will be suppressed by creating a custom function to obtain just the model estimates, thus avoiding repetition of the diagnostic plots. This will be achieved by calling the source code of gam.check and using the appropriate code in a new function that will be called gam_diagnostics, which can be found in the script gam_diagnostics.R.

In the next code chunk, we call gam.diagnostics to provide the desired diagnostic output:

```
source(here("scripts", "gam_diagnostics.R"))
gam_diagnostics(gam_00)
```

```
##
## Method: REML   Optimizer: outer newton
## full convergence after 5 iterations.
## Gradient range [-2.830761e-07,-3.314641e-08]
## (score 435.8783 & scale 388.2273).
## Hessian positive definite, eigenvalue range [0.009374965,49.00017].
## Model rank =  5 / 5
##
## Basis dimension (k) checking results. Low p-value (k-index<1) may
## indicate that k is too low, especially if edf is close to k'.
##
##          k'  edf k-index p-value
## s(Day) 4.00 1.18    0.35  <2e-16 ***
## ---
## Signif. codes:  0 '***' 0.001 '**' 0.01 '*' 0.05 '.' 0.1 ' ' 1
```

From the output of gam_diagnostics, it can be seen that the k-index is 0.35, which indicates that the model is not capturing the variability in the data. The edf (effective degrees of freedom) is an indicator of the complexity of the smooth. Here the complexity of the smooth is comparable to that of a 4th degree polynomial. And now obtain a summary of the fitted model, which is obtained by calling summary.

```
summary(gam_00)
```

```
##
## Family: gaussian
## Link function: identity
##
## Formula:
## StO2_sim ~ s(Day, k = 5)
##
## Parametric coefficients:
##             Estimate Std. Error t value Pr(>|t|)
## (Intercept)    20.17       1.97   10.24   <2e-16 ***
## ---
## Signif. codes:  0 '***' 0.001 '**' 0.01 '*' 0.05 '.' 0.1 ' ' 1
```

```
##
## Approximate significance of smooth terms:
##          edf Ref.df     F p-value
## s(Day) 1.184  1.342 8.861 0.00136 **
## ---
## Signif. codes:  0 '***' 0.001 '**' 0.01 '*' 0.05 '.' 0.1 ' ' 1
##
## R-sq.(adj) =  0.108   Deviance explained = 11.9%
## -REML = 435.88  Scale est. = 388.23    n = 100
```

From the `summary` function, information about the assumed distribution of the errors (Gaussian in this case) and the link function can be obtained. The link function is 'identity' as the model does not make any transformation on the predictors. The 'significance of smooth terms' *p-value* indicates if each smooth is significantly different from a constant mean 0 under the model. Here, the *p-value* is low but we have seen that there are issues with the model from the previous outputs. Finally, the 'deviance explained' indicates how much of the variation in the data the model is able to capture, which in this case corresponds to ≈ 12%.

### A.3 Second model

The major flaw of `gam_00` is that this model is not taking into account the fact that the data is nested in groups. The next iteration is a model where a different smooth of time (Day) is assigned for each group using `by = Group` in the model syntax. Below we fit such model, which is saved as `gam_01`, and use `gam_diagnostiscs` to obtain information from the model fit.

```
gam_01 <- gam(StO2_sim ~ s(Day, by = Group, k = 5), method = "REML", data
   = dat_sim)

gam_diagnostics(gam_01)
```

```
##
## Method: REML   Optimizer: outer newton
## full convergence after 9 iterations.
## Gradient range [-5.875578e-07,1.45076e-08]
## (score 400.4803 & scale 164.8662).
## Hessian positive definite, eigenvalue range [0.892525,48.57694].
## Model rank =  9 / 9
##
## Basis dimension (k) checking results. Low p-value (k-index<1) may
## indicate that k is too low, especially if edf is close to k'.
##
##                          k'  edf k-index p-value
## s(Day):GroupControl   4.00 3.69    0.58  <2e-16 ***
## s(Day):GroupTreatment 4.00 3.72    0.58  <2e-16 ***
## ---
## Signif. codes:  0 '***' 0.001 '**' 0.01 '*' 0.05 '.' 0.1 ' ' 1
```

Diagnostics for this model indicate that the k-index is still below 1 (0.58 from `gam_diagnostics` output), and that the residuals are still not following a normal distribution (Figure A.2). Moreover, the smooths (plotted via the `draw()` function) appear with a fairly linear profile, which indicates they are still not capturing the trends observed in the data.

```
summary(gam_01)
```

```
##
## Family: gaussian
## Link function: identity
##
## Formula:
## St02_sim ~ s(Day, by = Group, k = 5)
##
## Parametric coefficients:
##             Estimate Std. Error t value Pr(>|t|)
## (Intercept)   20.169      1.284   15.71   <2e-16 ***
## ---
## Signif. codes:  0 '***' 0.001 '**' 0.01 '*' 0.05 '.' 0.1 ' ' 1
##
## Approximate significance of smooth terms:
##                           edf Ref.df      F  p-value
## s(Day):GroupControl   3.688  3.942  9.084 2.33e-05 ***
## s(Day):GroupTreatment 3.723  3.954 33.465  < 2e-16 ***
## ---
## Signif. codes:  0 '***' 0.001 '**' 0.01 '*' 0.05 '.' 0.1 ' ' 1
##
## R-sq.(adj) =  0.621   Deviance explained =   65%
## -REML = 400.48  Scale est. = 164.87    n = 100
```

From summary(), the deviance explained by the model is ≈ 65%.

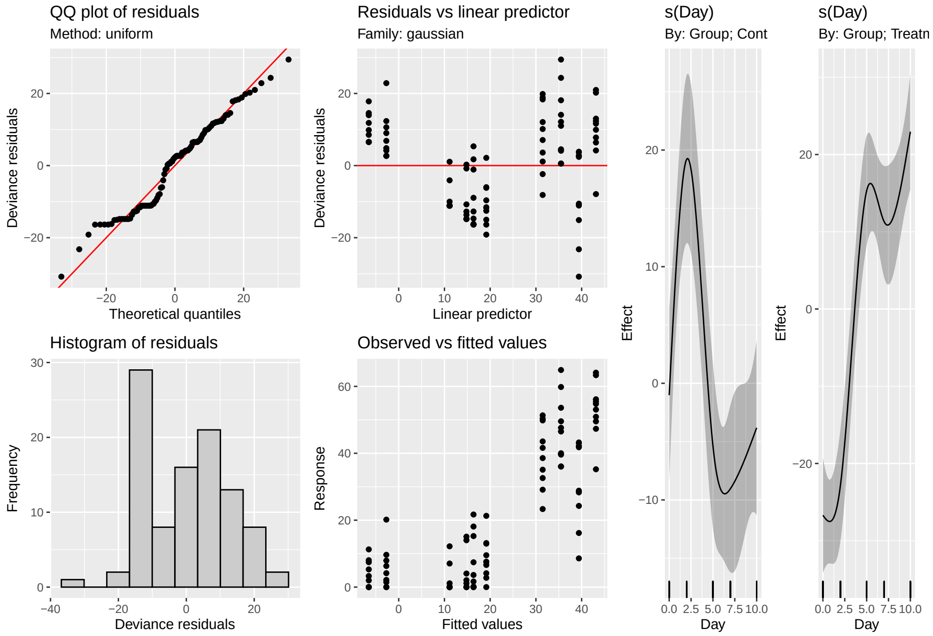


Figure A.2: Graphical diagnostics for the second GAM model. Left: Graphical diagnostics provided by the function appraise from the package *gratia*. Right: Fitted smooth for the model, provided by the function draw.

### A.4 Third model

Model gam_00 was built for didactic purposes to cover the simplest case, but it does not account for the nesting of the data by Group, which is apparent from the type of smooth fitted (a single smooth), the model diagnostics, and, the low variance explained by the model. On the other hand, gam_01 takes into account the nesting within each group and provides better variance explanation, but as indicated in Section 5.2 in the main manuscript, in order to differentiate between each group a parametric term needs to be added to the model for the interaction of *Day* and *Group*.

This is because in gam_01 separate smooths were fitted per group and those smooths also tried to account for the different means of the response in the two groups. Adding a parametric term for Group enables the smooths to capture the time course-differences of each group. The resulting model is gam_02, which is the model fitted in the main manuscript.

```
# GAM for StO2

gam_02 <- gam(StO2_sim ~ Group + s(Day, by = Group, k = 5), method = "REML
   ", data = dat_sim)

gam_diagnostics(gam_02)
```

```
##
## Method: REML   Optimizer: outer newton
## full convergence after 10 iterations.
## Gradient range [-2.934421e-07,2.346107e-07]
## (score 355.5265 & scale 63.48881).
## Hessian positive definite, eigenvalue range [1.21617,48.08942].
## Model rank =  10 / 10
##
## Basis dimension (k) checking results. Low p-value (k-index<1) may
## indicate that k is too low, especially if edf is close to k'.
##
##                          k'  edf k-index p-value
## s(Day):GroupControl   4.00 3.88    1.23    0.98
## s(Day):GroupTreatment 4.00 3.90    1.23    0.97
```

By using appraise() and draw on this model (Figure A.3 we see that the trend on the QQ plot has improved, the histogram of the residuals appears to be reasonably close to an normal distribution, and the smooths are capturing the trend of the data within each group. From gam_diagnostics, the k-index is now at an acceptable value ($\approx 1.23$).

```
summary(gam_02)
```

```
##
## Family: gaussian
## Link function: identity
##
## Formula:
## StO2_sim ~ Group + s(Day, by = Group, k = 5)
##
## Parametric coefficients:
##                Estimate Std. Error t value Pr(>|t|)
## (Intercept)      10.574      1.127   9.384 5.36e-15 ***
## GroupTreatment   19.191      1.594  12.043  < 2e-16 ***
## ---
## Signif. codes:  0 '***' 0.001 '**' 0.01 '*' 0.05 '.' 0.1 ' ' 1
##
## Approximate significance of smooth terms:
##                          edf Ref.df      F p-value
```

```
## s(Day):GroupControl    3.881   3.991 23.54   <2e-16 ***
## s(Day):GroupTreatment 3.904   3.994 87.29   <2e-16 ***
## ---
## Signif. codes:  0 '***' 0.001 '**' 0.01 '*' 0.05 '.' 0.1 ' ' 1
##
## R-sq.(adj) =  0.854   Deviance explained = 86.7%
## -REML = 355.53  Scale est. = 63.489     n = 100
```

From summary, the model is able to capture 86.7% of the variance in the data, which is a substantial improvement over the variance explained by gam_00 and gam_01.

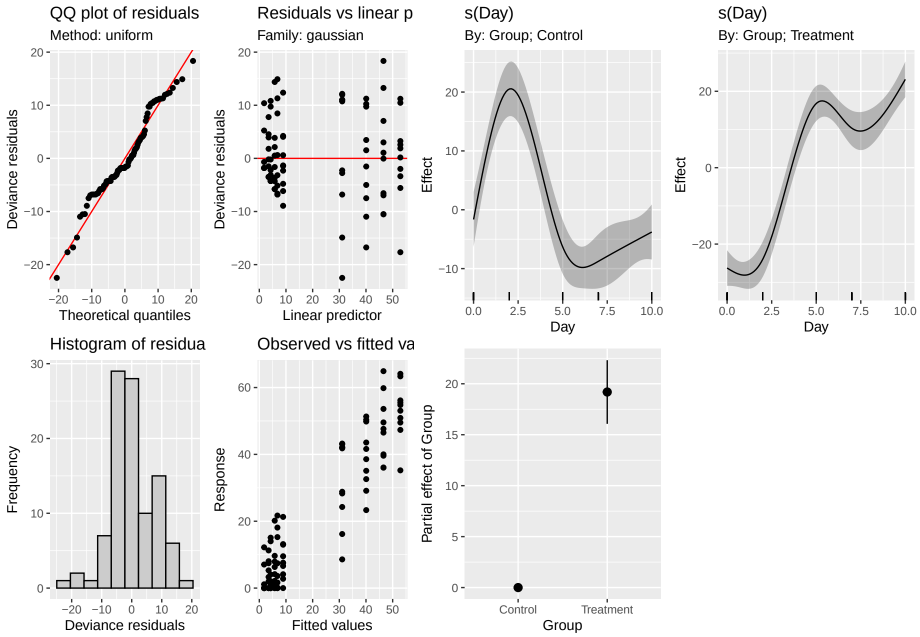


Figure A.3: Graphical diagnostics for the final GAM model. Left: Graphical diagnostics provided by the function appraise from the package *gratia*. Right: Fitted smooths for the model, provided by the function draw.

#### A.4.1 Comparing models via AIC

One final comparison that can be made for model selection involves the use of the Aikake Information Criterion (AIC). This metric is used to estimate information loss, which we want to minimize with an appropriate model. Therefore, when 2 or more models are compared, the model with lower AIC is preferred. In R, the comparison is done using the AIC function.

```
AIC(gam_00, gam_01, gam_02)
```

```
##               df      AIC
## gam_00  3.342420 884.4234
## gam_01  9.895821 805.3071
## gam_02 10.985458 710.5476
```

The output in this case is expected: model gam_02 has a lower AIC (710.54) whereas the initial two models have higher AICs (884.43 and 805.30). The AIC should not be considered as the only estimator of model quality, instead to be used as complimentary information to the graphical diagnostics and model checks described above.

### A.5 Pairwise comparisons of smooth confidence intervals

The estimation of significant differences between each treatment group can be achieved via pairwise comparisons of the smooth confidence intervals as described in Section 5.3 in the main manuscript.

In this case, the "design matrix" is used to estimate the pairwise comparisons (see main manuscript for details and associated references). Briefly, the "design matrix" (also known as the "Xp matrix") from the selected model (gam_02) is used to calculate a 95% confidence interval of the difference between the smooth terms for each group. This approach allows to estimate the time intervals where a significant difference exists between the groups (confidence interval above or below 0).

We want to emphasize that throught the manuscript and for the comparisons on gam_02, we included the group means in order to keep the pairwise comparisons on the scale of the response. By default, pairwise comparisons available in other software packages (such as in G. Simpson's *gratia*) do not include the group means. However, we decided to include them because for our approach, it is easier to see the magnitude in the change of the difference between two treatment groups when the means are included.

The change we allude to (mean group inclusion) can be found in the script pointwise_comparisons.R, at the line where we have added a comment that reads:

```
###### IMPORTANT: uncommenting the following two lines removes the group
   means
###### from the comparison######
```

The inclusion of group means works well for models like gam_02, but implementing this on models with more parametric terms can be challenging and we want the reader to be aware of this.

However, we do believe that the model presented in the paper covers a wide range of situations and our approach here for the pairwise comparisons will be useful for most biomedical researchers. We first call the script pointwise_comparisons.R to estimate the pointwise difference, then call the script difference_smooths.R which calculates the simultaneous confidence interval for the difference, and finally pairwise_limits.R, which estimates the regions to be shaded for the intervals with significance in the final figure. We use difference_smooths on model gam_02 and save the estimates in diff_complete.

```
source(here::here("scripts", "pointwise_comparisons.R"))
source(here::here("scripts", "difference_smooths.R"))
source(here::here("scripts", "pairwise_limits.R"))
# compute difference between smooths and calculate confidence interval:
# complete data
diff_complete <- difference_smooths(gam_02, smooth = "s(Day)", newdata =
   newdat,
    unconditional = TRUE, frequentist = FALSE, n = 100, partial_match =
       TRUE, nrep = 10000)
my_list <- pairwise_limits(diff_complete)
```

Next, we plot the comparisons from diff_complete and create a ggplot2 object (c1) so we can also shade the regions of significant differences easily.

```
rib_col <- "#8D7D82"  #color for ribbon for confidence interval
control_rib <- "#875F79"  #color for ribbon for control region
treat_rib <- "#A7D89E"  #color for ribbon treatment region

c1 <- ggplot() + geom_line(data = diff_complete, aes(x = Day, y = diff),
   size = 1,
    alpha = 0.5) + annotate("rect", xmin = my_list$init1, xmax = my_list$
       final1,
    ymin = -Inf, ymax = Inf, fill = control_rib, alpha = 0.5) + annotate("
       text",
```

```
    x = 1.5, y = -18, label = "Control>Treatment", size = 6, angle = 90) +
     annotate("rect",
    xmin = my_list$init2, xmax = my_list$final2, ymin = -Inf, ymax = Inf,
       fill = treat_rib,
    alpha = 0.5) + annotate("text", x = 6, y = -18, label = "Treatment>
       Control",
    size = 6, angle = 90) + geom_ribbon(data = diff_complete, aes(x = Day,
       ymin = lower_s,
    ymax = upper_s), alpha = 0.5, fill = rib_col, inherit.aes = FALSE) +
       geom_hline(yintercept = 0,
    lty = 2, color = "red") + scale_x_continuous(breaks = c(0, 2, 5, 7,
       10)) + labs(y = "Difference\n(Complete observations)") +
    theme_classic() + theme(axis.text = element_text(size = 22))
```

The resulting plot (Figure A.4) shows the estimated difference in the smooths with a 95% simultaneous CI. As a reminder, because we have kept the group means, we can directly see that the Treatment starts to have a significant effect around day 3. As therapy progresses, the effect continues, and the magnitude of the difference (≈ 40%) at day five corresponds directly with the magnitude in the increase in $StO_2$ in the group at the same time point in the data.

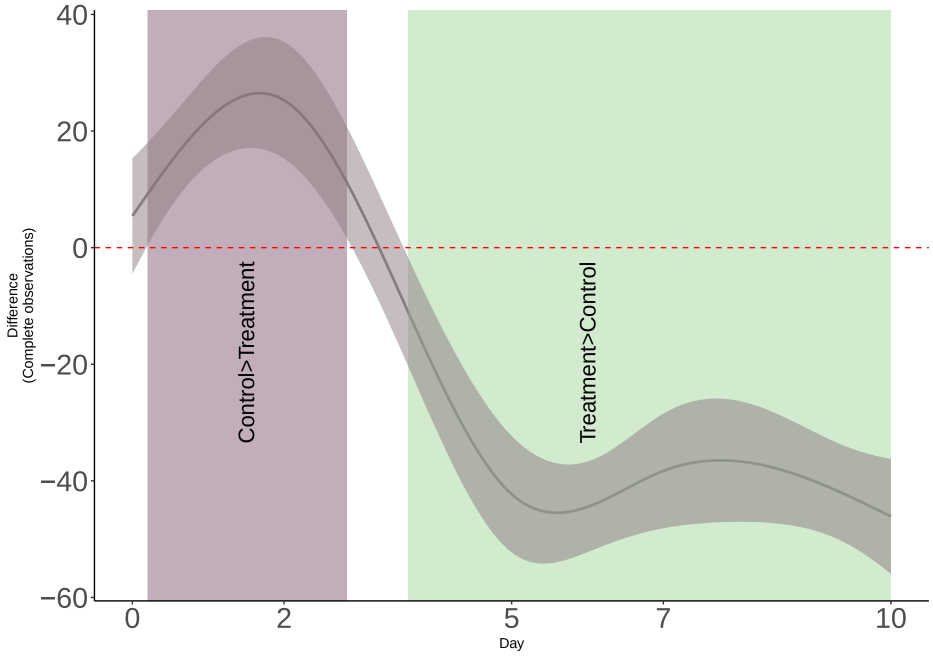


Figure A.4: Smooth pairwise comparisons for model `gam_02` using a 95% simultaneous CI for the difference between smooths. The comparison includes the group means and therefore can be directly correlated with the magnitude of the response. Shaded regions indicate time intervals where each treatment group has a non-zero effect.

### A.6 Final considerations

In this Appendix we have shown a basic model selection workflow for GAMs. Our goal here is to familiarize the reader with the logic behind the construction of each model and the kind of diagnostic information that needs to be checked to ensure the model is adequate. One final consideration must be given to the conditional distribution of the response. In the main manuscript and through this workflow, we have assumed a normal distribution for the response, which we believe is appropriate in many biomedical scenarios. However, we must remind the reader that the normal distribution is not intended to be used as a "one size fits all" distribution.

Depending on the type of data (counts, binary outcomes) the user can choose different conditional distributions in *mgcv*.

We also want to indicate that one of the major advantages of fitting GAMs in R using *mgcv* is that once the appropriate conditional distribution has been chosen and model diagnostics are assessed, the user only needs to choose a basis dimension (the number of basis functions to use), and check if the basis dimension is adequately capturing the trend of the data using `k.check`. If that is not the case, then the user needs to increase `k` a bit and check `k.check` again. The process might seem mechanistic, but we hope that with the theory presented in the main manuscript and the workflow of this Appendix the user has a good understanding of what the model is doing and the rationale for choosing a GAM as a tool for statistical analysis.

### A.7 Additional resources

Multiple and excellent resources are available for biomedical researchers that want to gain more insight on the theory and computation of GAMs. Here we provide a brief list of resources that cover additional topics as well as available packages that are worth considering:

- Gavin Simpson's package *gratia*, which provides convenient wrapper functions for plotting and pairwise comparisons. CRAN page: https://cran.r-project.org/web/packages/gratia/gratia.pdf

Vignette:

https://gavinsimpson.github.io/gratia/

- Gavin Simpson's blog *From the bottom of the heap*, which covers a wide range of topics in GAM modeling, news about updates on *gratia* and provides tutorials on GAM fitting.

Link:

https://fromthebottomoftheheap.net/

- Matteo Fasiolo's package *mgcViz*, ane extension of the *mgcv* package. Provides visual tools for Generalized Additive Models that exploit the additive structure of such models.

CRAN Page:

https://cran.r-project.org/web/packages/mgcViz/index.html

Vignette: https://mfasiolo.github.io/mgcViz/articles/mgcviz.html

- The book "Generalized Additive Models: An Introduction with R" (the 2nd. Edition can be found here) by Simon Wood is an excellent resource for in-depth material on GAM theory, mathematical derivation, examples, and detailed descriptions on the computational aspect of *mgcv*.

Finally, Noam Ross has assembled a list of online tools and resources for learning about and using GAMs in R, which cover a range of very useful videos, slides, and courses from different authors. The list can be found in Noam Ross's GitHub profile here:

Resources for Learning About and Using GAMs in R
