## Appendix B: Code for "Generalized additive models to analyze non-linear trends in biomedical longitudinal data using R: Beyond repeated measures ANOVA and Linear Mixed Models"

Ariel I. Mundo, John R. Tipton, and Timothy J. Muldoon*

**

This appendix shows the code for the functions used through the main manuscript, which can be found in the *scripts* folder in the GitHub repository. We provide a brief explanation of the purpose of each function.

### B.1 Setup

First, we load all required libraries and set seed.

```
library(patchwork)
library(tidyverse)
library(mvnfast)
library(nlme)
library(mgcv)
library(gratia)
library(here)
library(scico)
set.seed(2021)  #set seed for reproducibility

# alpha for ribbon in the smooth plots
al <- 0.8

thm1 <- scale_fill_scico_d(palette = "tokyo", begin = 0.3, end = 0.8,
   direction = -1,
    aesthetics = c("colour", "fill"))
```

### B.2 Linear and quadratic longitudinal trends

#### B.2.1 Function for linear and quadratic trends, rm-ANOVA and LMEM fits

The first function is `example()` which is in the file named `example.R` in the `scripts/` folder, which allows to simulate linear and quadratic data in the same manner as in Section 3.5 in the main manuscript. Both rm-ANOVA and LMEM with interaction are fitted to the data. The error for each simulated trend can be correlated or uncorrelated as well.

```
##########Section for calculations###########

## Example with linear response

#This function simulates data using a linear or quadratic mean response
   and each with correlated
#or uncorrelated errors. Each group has a different slope/concavity.
example <- function(n_time = 6, #number of time points
                    fun_type = "linear", #type of response
                    error_type = "correlated") {

  if (!(fun_type %in% c("linear", "quadratic")))
    stop('fun_type must be either "linear", or "quadratic"')
  if (!(error_type %in% c("correlated", "independent")))
    stop('fun_type must be either "correlated", or "independent"')


  x <- seq(1,6, length.out = n_time)

  #Create mean response matrix: linear or quadratic
  mu <- matrix(0, length(x), 2)
```

```r
if (fun_type == "linear") {
  # linear response
  mu[, 1] <- -(0.25*x)+2
  mu[, 2] <- 0.25*x+2
} else {
  # quadratic response (non-linear)
  mu[, 1] <- -(0.25 * x^2) + 1.5 * x - 1.25
  mu[, 2] <- (0.25 * x^2) - 1.5 * x + 1.25
}


#create an array where individual observations per each time point for
   each group are to be stored. Currently using 10 observations per
   timepoint
y <- array(0, dim = c(length(x), 2, 10))

#Create array to store the "errors" for each group at each timepoint.
   The "errors" are the between-group variability in the response.
errors <- array(0, dim = c(length(x), 2, 10))
#create an array where 10 observations per each time point for each
   group are to be stored

#The following loops create independent or correlated responses. To each
    value of mu (mean response per group) a randomly generated error (
   correlated or uncorrelated) is added and thus the individual response
    is created.
if (error_type == "independent") {
  ## independent errors
  for (i in 1:2) {
    for (j in 1:10) {
      errors[, i, j] <- rnorm(6, 0, 0.25)
      y[, i, j] <- mu[, i] + errors[, i, j]
    }
  }
} else {
  for (i in 1:2) {     # number of treatments
    for (j in 1:10) {  # number of subjects
      # compound symmetry errors: variance covariance matrix
      errors[, i, j] <- rmvn(1, rep(0, length(x)), 0.1 * diag(6) + 0.25
         * matrix(1, 6, 6))
      y[, i, j] <- mu[, i] + errors[, i, j]
    }
  }
}


## subject random effects

## visualizing the difference between independent errors and compound
   symmetry
## why do we need to account for this -- overly confident inference

#labeling y and errors
dimnames(y) <- list(time = x, treatment = 1:2, subject = 1:10)
```

```r
  dimnames(errors) <- list(time = x, treatment = 1:2, subject = 1:10)

  #labeling the mean response
  dimnames(mu) <- list(time = x, treatment = 1:2)

  #convert y, mu and errors to  dataframes with time, treatment and
     subject columns
  dat <- as.data.frame.table(y, responseName = "y")
  dat_errors <- as.data.frame.table(errors, responseName = "errors")
  dat_mu <- as.data.frame.table(mu, responseName = "mu")

  #join the dataframes to show mean response and errors per subject
  dat <- left_join(dat, dat_errors, by = c("time", "treatment", "subject")
     )
  dat <- left_join(dat, dat_mu, by = c("time", "treatment"))
  #add time
  dat$time <- as.numeric(as.character(dat$time))
  #label subjects per group
  dat <- dat %>%
    mutate(subject = factor(paste(subject,
                                  treatment,
                                  sep = "-")))


  ## repeated measures ANOVA

  fit_anova <- lm(y ~ time + treatment + time * treatment, data = dat)

  #LMEM: time and treatment interaction model, compound symmetry
  fit_lme <- lme(y ~ treatment + time + treatment:time,
                 data = dat,
                 random = ~ 1 | subject,
                 correlation = corCompSymm(form = ~ 1 | subject))

  #create a prediction frame where the model can be used for plotting
     purposes
  pred_dat <- expand.grid(treatment = factor(1:2),
                          time = unique(dat$time))

  #add model predictions to the dataframe that has the simulated data
  dat$pred_anova <- predict(fit_anova)
  dat$pred_lmem <- predict(fit_lme)

  #return everything in a list
  return(list(
    dat       = dat,
    pred_dat  = pred_dat,
    fit_anova = fit_anova,
    fit_lme   = fit_lme
  ))
}
```

#### B.2.2 A composite plot for the trends

The function `plot_example()` from the file `plot_example.R` in the folder `scripts/` uses the output of `example.R` to show the fit of a rm-ANOVA and a LMEM. This function can be used to show an expanded version of Figure 1 in the main manuscript, presenting simulated data with correlated and uncorrelated errors and how the individual trends vary in each case. The corresponding rm-ANOVA and LMEM fits are also presented, and we show the complete output in the next subsection.

```
## This function plots the rm-ANOVA and LMEM for the data simulated in
## example.R
plot_example <- function(sim_dat) {
    txt <- 20
    p1 <- sim_dat$dat %>%
        ggplot(aes(x = time, y = y, group = treatment, color = treatment))
            + geom_point(show.legend = FALSE) +
        labs(y = "response") + geom_line(aes(x = time, y = mu, color =
           treatment),
        show.legend = FALSE) + theme_classic() + theme(plot.title =
           element_text(size = txt,
        face = "bold"), text = element_text(size = txt)) + thm1

    # plot the simulated data with trajectories per each subject
    p2 <- sim_dat$dat %>%
        ggplot(aes(x = time, y = y, group = subject, color = treatment)) +
            geom_line(aes(size = "Subjects"),
        show.legend = FALSE) + # facet_wrap(~ treatment) + show.legend =
           FALSE)
        show.legend = FALSE) + # facet_wrap(~ treatment) + + # facet_wrap(
           ~
        show.legend = FALSE) + # facet_wrap(~ treatment) + treatment) +
    geom_line(aes(x = time, y = mu, color = treatment, size = "Simulated
       Truth"),
        lty = 1, show.legend = FALSE) + labs(y = "response") + scale_size_
           manual(name = "Type",
        values = c(Subjects = 0.5, `Simulated Truth` = 3)) + theme_classic
           () + theme(plot.title = element_text(size = txt,
        face = "bold"), text = element_text(size = txt)) + thm1

    # plot the errors
    p3 <- sim_dat$dat %>%
        ggplot(aes(x = time, y = errors, group = subject, color =
           treatment)) + geom_line(show.legend = FALSE) +
        labs(y = "errors") + theme_classic() + theme(plot.title = element_
           text(size = txt,
        face = "bold"), text = element_text(size = txt)) + thm1

    # plot the model predictions for rm-ANOVA
    p4 <- ggplot(sim_dat$dat, aes(x = time, y = y, color = treatment)) +
       geom_point(show.legend = FALSE) +
        labs(y = "response") + geom_line(aes(y = predict(sim_dat$fit_anova
           ), group = subject,
        size = "Subjects"), show.legend = FALSE) + geom_line(data = sim_
           dat$pred_dat,
        aes(y = predict(sim_dat$fit_anova, level = 0, newdata = sim_dat$
           pred_dat),
```

```
            size = "Population"), show.legend = FALSE) + guides(color =
               guide_legend(override.aes = list(size = 2))) +
        scale_size_manual(name = "Predictions", values = c(Subjects = 0.5,
            Population = 3)) +
        theme_classic() + theme(plot.title = element_text(size = txt, face
            = "bold"),
        text = element_text(size = txt)) + thm1

    # plot the LMEM predictions
    p5 <- ggplot(sim_dat$dat, aes(x = time, y = y, color = treatment)) +
       geom_point() +
        labs(y = "response") + geom_line(aes(y = predict(sim_dat$fit_lme),
            group = subject,
        size = "Subjects")) + geom_line(data = sim_dat$pred_dat, aes(y =
           predict(sim_dat$fit_lme,
        level = 0, newdata = sim_dat$pred_dat), size = "Population")) +
           guides(color = guide_legend(override.aes = list(size = 2))) +
        scale_size_manual(name = "Predictions", values = c(Subjects = 0.5,
            Population = 3)) +
        theme_classic() + theme(plot.title = element_text(size = txt, face
            = "bold"),
        text = element_text(size = txt)) + thm1

    if (option == "simple") {
        return((p1 + p4 + p5) + plot_layout(nrow = 1) + plot_annotation(
           tag_levels = "A"))
    } else {
        return((p1 + p3 + p2 + p4 + p5) + plot_layout(nrow = 1) + plot_
           annotation(tag_levels = "A"))
    }
}
```

#### B.2.3 Plotting rm-ANOVA and LMEM fits for linear and quadratic trends in data

In this subsection, we use the `example()` and `plot_example()` functions to create an expanded version of Figure 1 in the main manuscript. The difference here is that in the main manuscript `plot_example()` uses `option='simple'` to create the plot, and therefore only shows the simulated data, and the rm-ANOVA and LMEM fit to the simulated data with correlated errors. Here, we use the option `composite` to generate the additional panels for the uncorrelated errors and their respective rm-ANOVA and LMEM fits.

Figure B.1 show in panels A and D the simulated mean responses and individual data points. Panels C and G show a visual interpretation of “correlation” in the responses: In panel C, subjects that have a value of the random error $\varepsilon$ either above or below the mean group response are more likely to have other observations that follow the same trajectory, thereby demonstrating correlation in the response. In panel G,because the errors are independent, there is no expectation that responses are likely to follow a similar pattern. Panels D and H show the predictions from the rm-ANOVA model.

**B.2.3.1 Fits for linear trends** The chunk below sources both `example.R` and `plot_example_Appendix.R` files to simulate data and create the composite plots.

```
source(here::here("scripts", "example.R"))
source(here::here("scripts", "plot_example.R"))

A1 <- plot_example(example(fun_type = "linear", error_type = "correlated")
   , option = "composite")
```

```
B1 <- plot_example(example(fun_type = "linear", error_type = "independent"
   ), option = "composite")

C1 <- plot_example(example(fun_type = "quadratic", error_type = "
   correlated"), option = "composite")

D1 <- plot_example(example(fun_type = "quadratic", error_type = "
   independent"), option = "composite")
```

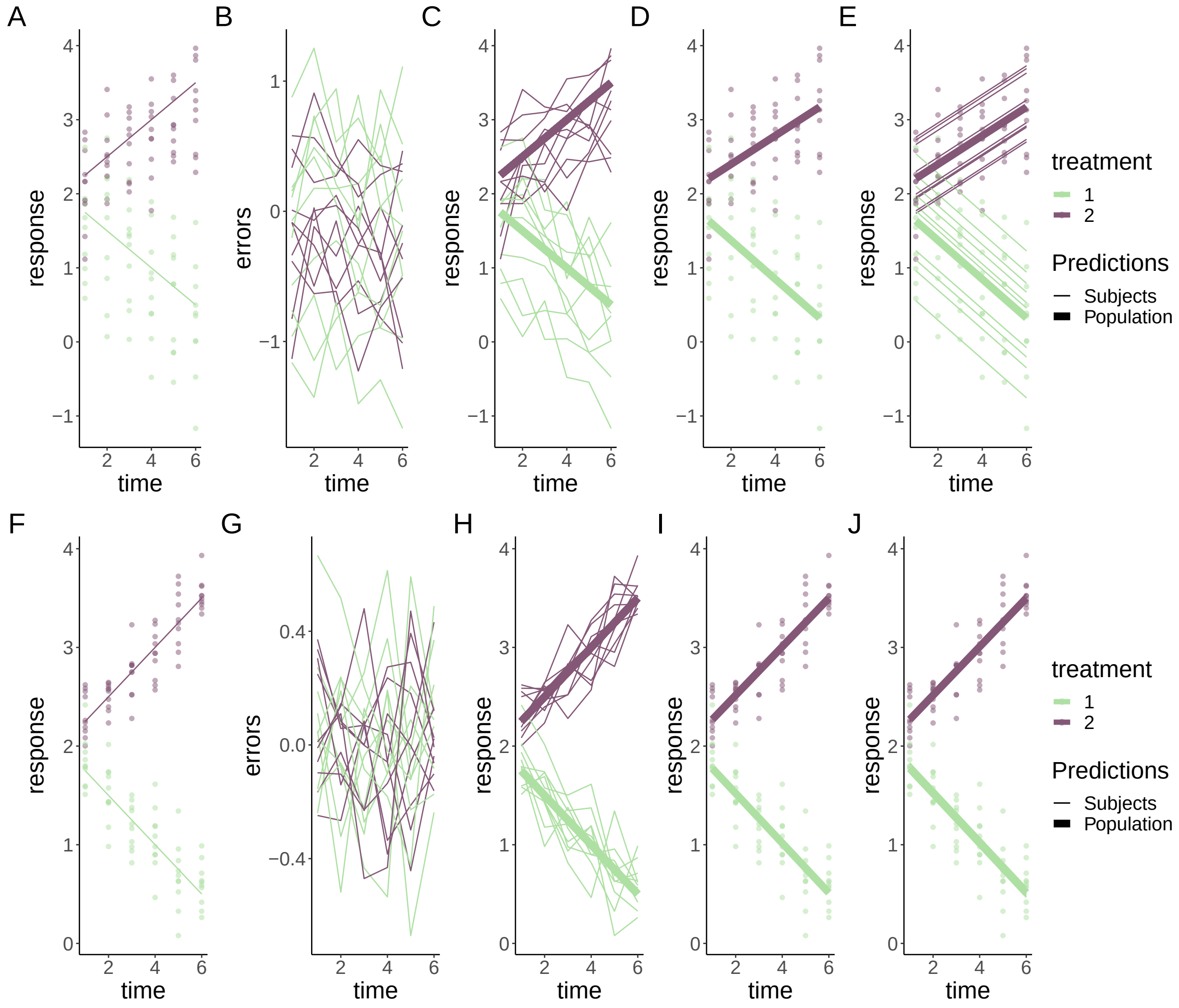


Figure B.1: Simulated linear responses from two groups with correlated (top row) or independent (bottom row) errors using a rm-ANOVA model and a LMEM. **A, F**:Simulated data with known mean response and individual responses (points) showing the dispersion of the data. **B, G**: Generated errors showing the difference in the behavior of correlated and independent errors. **C, H**: Simulated data with thin lines representing individual trajectories. **D, I**: Estimations from the rm-ANOVA model for the mean group response. **E, J**: Estimations from the LMEM for the mean group response and individual responses (thin lines). In all panels, thick lines are the predicted mean response per group, thin lines are the random effects for each subject and points represent the original raw data. Both rm-ANOVA and the LMEM are able to capture the trend of the data.

**B.2.3.2 Fits for quadratic trends** For the quadratic response case, Figure B.2 shows the simulated responses using compound symmetry and independent errors.

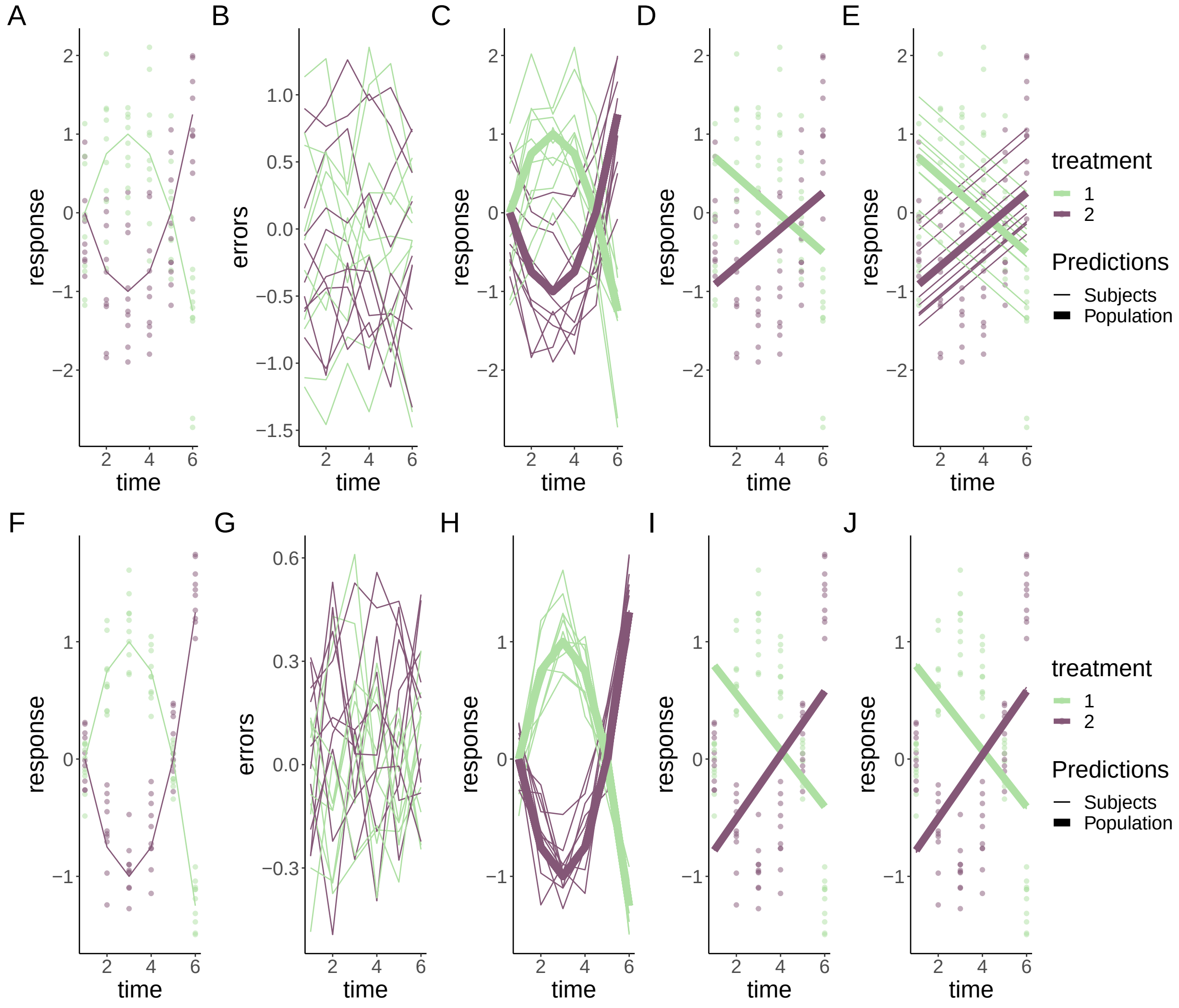


Figure B.2: Simulated quadratic responses from two groups with correlated (top row) or independent (bottom row) errors using a rm-ANOVA model and a LMEM. **A, F**:Simulated data with known mean response and individual responses (points) showing the dispersion of the data. **B, G**: Generated errors showing the difference in the behavior of correlated and independent errors. **C, H**: Simulated data with thin lines representing individual trajectories. **D, I**: Estimations from the rm-ANOVA model for the mean group response. **E, J**: Estimations from the LMEM for the mean group response and individual responses (thin lines). In all panels, thick lines are the predicted mean response per group, thin lines are the random effects for each subject and points represent the original raw data. Both rm-ANOVA and the LMEM are not able to capture the changes in each group over time.

### B.3 Basis Functions

The next chunk presents the code used to create Figure 2 in the main manuscript. Here, we simulate data from the same concave down quadratic trend from Figure B.2A and use the package *mgcv* to fit a GAM to fit the trend of the data over time. The GAM in this case can be represented as

$$Response = \beta_0 + f(time_t|\beta_1)$$

Because we only use the data from one group and we do not account for interaction. The model in *mgcv* is written as:

```
gm<-gam( y~s(time,k = 5),data = dat, method = "REML")
```

From the model, we extract the original and weighted four basis functions (and the intercept), and add these last weighted basis functions to construct the smoother.

```
#basis functions: this script creates Fig 2. by calculating a GAM for the
   Group 1 data of Figure 1,
# extracting the basis functions
#and creates objects p11,p12,p13,p14 for plotting, which are combined in b
   _plot to create the
#final composite figure
n_time = 6
x <- seq(1,6, length.out = n_time)
mu <- matrix(0, length(x), 2)
mu[, 1] <- -(0.25 * x^2) + 1.5 * x - 1.25 #mean response
mu[, 2] <- (0.25 * x^2) - 1.5 * x + 1.25 #mean response
y <- array(0, dim = c(length(x), 2, 10))
errors <- array(0, dim = c(length(x), 2, 10))
for (i in 1:2) {     # number of treatments
  for (j in 1:10) {  # number of subjects
    # compound symmetry errors
    errors[, i, j] <- rmvn(1, rep(0, length(x)), 0.1 * diag(6) + 0.25 *
       matrix(1, 6, 6))
    y[, i, j] <- mu[, i] + errors[, i, j]
  }
}

#label each table
dimnames(y) <- list(time = x, treatment = 1:2, subject = 1:10)
dimnames(errors) <- list(time = x, treatment = 1:2, subject = 1:10)
dimnames(mu) <- list(time = x, treatment = 1:2)

#Convert to dataframes with subject, time and group columns
dat <- as.data.frame.table(y, responseName = "y")
dat_errors <- as.data.frame.table(errors, responseName = "errors")
dat_mu <- as.data.frame.table(mu, responseName = "mu")
dat <- left_join(dat, dat_errors, by = c("time", "treatment", "subject"))
dat <- left_join(dat, dat_mu, by = c("time", "treatment"))
dat$time <- as.numeric(as.character(dat$time))

#label subject per group
dat <- dat %>%
  mutate(subject = factor(paste(subject, treatment, sep = "-")))

#extract  "Group 1" to fit the GAM
dat <- subset(dat, treatment==1)
#keep just the response and timepoint columns
dat <- dat[,c('y', 'time')]

#GAM model of time, 5 basis functions
gm <- gam(y ~ s(time, k = 5), data = dat, method = "REML")
```

```
#model_matrix (also known as) 'design matrix'
#will contain the smooths used to create  model 'gm'
model_matrix <- as.data.frame(predict(gm, type = 'lpmatrix'))


time <- c(1:6)

basis <- model_matrix[1:6, ] #extracting basis (because the values are
   repeated after every 6 rows)
#basis<-model_matrix[1:6,-1] #extracting basis
colnames(basis)[colnames(basis) == "(Intercept)"] <- "s(time).0"
basis <- basis %>% #pivoting to long format
  pivot_longer(cols = starts_with("s")) %>%
  arrange(name) #ordering

#length of dataframe to be created: number of basis by number of
   timepoints (minus 1 for the intercept that we won't plot)
ln <- 6 * (length(coef(gm)))

basis_plot <- data.frame(Basis = integer(ln),
                         value_orig = double(ln),
                         time = integer(ln),
                         coef = double(ln))

basis_plot$time <- rep(time) #pasting timepoints
basis_plot$Basis <- factor(rep(c(1:5), each = 6)) #pasting basis number
   values
basis_plot$value_orig <- basis$value #pasting basis values
basis_plot$coef <- rep(coef(gm)[1:5], each = 6) #pasting coefficients
basis_plot <- basis_plot %>%
  mutate(mod_val = value_orig * coef) #the create the predicted values the
      bases need to be
#multiplied by the coefficients

#creating labeller to change the labels in the basis plots

basis_names <- c(`1`="Intercept", `2`="1", `3`="2", `4`="3", `5`="4")

#calculating the final smooth by aggregating the basis functions

smooth <- basis_plot %>%
  group_by(time) %>%
  summarize(smooth = sum(mod_val))


#original basis
sz <- 1
p11 <- ggplot(basis_plot,
              aes(x = time, y = value_orig, colour = as.factor(Basis))) +
  geom_line(size=sz, show.legend = FALSE) +
  geom_point(size = sz + 1, show.legend = FALSE) +
  labs(y = 'Basis functions') +
  facet_wrap(~ Basis, labeller = as_labeller(basis_names)) +
  theme_classic() +
```

```
  thm1

#penalized basis
p12 <- ggplot(basis_plot,
              aes(x = time, y = mod_val, colour = as.factor(Basis))) +
  geom_line(show.legend = FALSE, size = sz) +
  geom_point(show.legend = FALSE, size = sz + 1) +
  labs(y = 'Penalized \n basis functions') +
  scale_y_continuous(breaks = seq(-1, 1, 1)) +
  facet_wrap(~ Basis, labeller = as_labeller(basis_names)) +
  theme_classic() +
  thm1

#heatmap of the  coefficients
x_labels <- c("Intercept", "1", "2", "3", "4")
p13 <- ggplot(basis_plot,
              aes(x = Basis, y = Basis)) +
  geom_tile(aes(fill = coef), colour = "black") +
  scale_fill_gradient(low = "white", high = "#B50A2AFF") +
  labs(x = 'Basis', y = 'Basis') +
  scale_x_discrete(labels = x_labels) +
  geom_text(aes(label = round(coef, 2)),
            size = 7, show.legend = FALSE) +
  theme_classic()+
  theme(legend.title = element_blank())

#plotting simulated datapoints and smooth term
p14<-ggplot(data = dat,
            aes(x = time, y = y)) +
  geom_point(size = sz + 1, alpha = 0.5) +
  thm1 +
  labs(y = 'Simulated \n response') +
  geom_line(data = smooth, aes(x = time, y = smooth),
            color = "#6C581DFF", size = sz + 1)+
  theme_classic()


#Combining all
b_plot <- p11 + p13 + p12 + p14 + plot_annotation(tag_levels = 'A') &
  theme(text = element_text(size=18))
```

### B.4 Function for data simulation

The next chunk presents the code of the function `simulate_data()` from the file `simulate_data.R` in the `scripts/` folder, which we use in the main manuscript to simulate data that follows the reported trends in $StO_2$ in the literature. Although the default standard deviation (SD) to generate the normally-distributed data in the function is 10%, we use a standard deviation of 10% in the main manuscript. We use the default number of 10 observations at each time point in the simulated data. Note that the input for the function is `dat`, needs to be a dataframe that contains the original values for the trends in each group over time from the literature. In our manuscript, we use the dataframe

```
# data from Vishwanath et. al.
dat <- tibble(StO2 = c(4, 27, 3, 2, 0.5, 7, 4, 50, 45, 56), Day = rep(c(0,
    2, 5,
```

```
    7, 10), times = 2), Group = as.factor(rep(c("Control", "Treatment"),
        each = 5)))
```

As the input for the function. Here, dataframe contains the trends in $StO_2$ for each group through the five time points and the labels for each treatment group (`Control` or `Treatment`).

```
# This function simulates data for the tumor data using default parameters
    of
# 10 observations per time point,and Standard deviation (sd) of 5%.
   Because
# physiologically StO2 cannot go below 0%, data is generated with a cutoff
# value of 0.0001 (the 'StO2_sim')


simulate_data <- function(dat, n = 10, sd = 5) {
    dat_sim <- dat %>%
        slice(rep(1:n(), each = n)) %>%
        group_by(Group, Day) %>%
        mutate(StO2_sim = pmax(rnorm(n, StO2, sd), 1e-04), subject = rep
            (1:10), subject = factor(paste(subject,
             Group, sep = "-"))) %>%
        ungroup()

    return(dat_sim)
}
```

### B.5 Pointwise and simultaneous confidence intervals for smooths

In this subsection we present the code used to create the pointwise and simultaneous empirical Bayesian confidence intervals (CIs) that are shown along the fitted smooths for the data in Figure 3 in the main manuscript. The computation is the same in the case of the complete and incomplete simulated data, and therefore we present here only the code for the incomplete data case.

We do not use a custom function in this case but instead extract the pointwise and simultaneous CIs after fitting the GAM by using the function `confint` methods from the *gratia* package. The boundaries for the pairwise and simultaneous CIs are obtained by using the type `"confindence"` or `"simultaneous"` in `confint` and are stored in `ci` and `si` respectively.

Because we want to include the group means for plotting purposes, we extract the intercept (`const`) and add it to the Treatment group estimates in both `ci` and `si` order to shift the interval so it matches the scale of the response. Finally, we create a *ggplot2* object for plotting.

```
# the same model used for the full dataset
mod_m1 <- gam(StO2_sim ~ Group + s(Day, by = Group, k = 5), data = dat_
   missing)
# appraise the model
appraise(mod_m1)


ci <- confint(mod_m1, parm = "s(Day)", partial_match = TRUE, type = "
   confidence")
## simultaneous interval
si <- confint(mod_m1, parm = "s(Day)", type = "simultaneous", partial_
   match = TRUE)
```

```
# mean shift for group 2
const <- coef(mod_m1)[2]

ci <- ci %>%
    mutate(est = case_when(Group == "Treatment" ~ est + const, TRUE ~ est)
        , lower = case_when(Group ==
        "Treatment" ~ lower + const, TRUE ~ lower), upper = case_when(
           Group == "Treatment" ~
        upper + const, TRUE ~ upper))

si <- si %>%
    mutate(est = case_when(Group == "Treatment" ~ est + const, TRUE ~ est)
        , lower = case_when(Group ==
        "Treatment" ~ lower + const, TRUE ~ lower), upper = case_when(
           Group == "Treatment" ~
        upper + const, TRUE ~ upper))


f6 <- ggplot(ci, aes(x = Day, y = est, group = smooth)) + geom_line(lwd =
   1) + geom_ribbon(data = ci,
    aes(ymin = lower, ymax = upper, x = Day, group = smooth, fill = Group)
       , inherit.aes = FALSE,
    alpha = 0.7, show.legend = FALSE) + geom_ribbon(data = si, aes(ymin =
       lower,
    ymax = upper, x = Day, group = smooth, fill = Group), inherit.aes =
       FALSE, alpha = 0.4,
    show.legend = TRUE) + geom_line(data = si, aes(Day, upper, color =
       Group), size = 0.8,
    alpha = 0.7) + geom_point(data = dat_missing, aes(x = Day, y = StO2_
       sim, color = Group),
    size = 1.5, alpha = 0.6, inherit.aes = FALSE) + labs(y = expression(
       atop(StO[2],
    "(incomplete observations)"))) + scale_x_continuous(breaks = c(0, 2,
       5, 7, 10)) +
    theme_classic() + theme(axis.text = element_text(size = 22)) + thm1
```

### B.6 Pairwise comparisons

This subsection presents the code used in functions `pointwise_comparisons()` and `difference_smooths()` from the files `pointwise_comparisons.R` and `difference_smooths.R` in the `scripts/` folder, which generate the across the function (pointwise) and simultaenous CI for the estimated difference between the smooths. In essence, `pointwise_comparisons()` computes the difference between the smooths and the pointwise CI. Then, `difference_comparisons()` uses the output of `pointwise_comparisons.R` to estimate the simultaneous CI.

The chunk below presents the code for `pointwise_comparisons.R`. Note that as we indicate in Appendix A in this case we include the group means to estimate the confidence intervals so they can be directly compared with the original data. This works for the model we used in the paper, but if there are more parametric terms in the model the computation is not straightforward and will require a careful implementation because with an increase in parametric terms the number of pairwise comparisons also increases greatly.

```
##this function determines the pointwise confidence interval of a
   difference between two smooths
```

```
difference_pointwise <- function(f1, f2, smooth, by_var, smooth_var,
                                 data, Xp, V, coefs, nrep = 1000) {
  ## make sure f1 and f2 are characters
  f1 <-  as.character(f1)
  f2 <-  as.character(f2)
  cnames <- colnames(Xp)
  ## columns of Xp associated with pair of smooths
  c1 <- grepl(gratia:::mgcv_by_smooth_labels(smooth, by_var, f1),
              cnames, fixed = TRUE)
  c2 <- grepl(gratia:::mgcv_by_smooth_labels(smooth, by_var, f2),
              cnames, fixed = TRUE)
  ## rows of Xp associated with pair of smooths
  r1 <- data[[by_var]] == f1
  r2 <- data[[by_var]] == f2

  ## difference rows of Xp for pair of smooths
  X <- Xp[r1, ] - Xp[r2, ]

  ######IMPORTANT: uncommenting the following two lines
  #removes the group means from the comparison######

  ## zero the cols related to other splines
  # X[, ! (c1 | c2)] <- 0

  ## zero out the parametric cols
  #X[, !grepl('^s\\(', cnames)] <- 0

  ## compute difference
  sm_diff <- drop(X %*% coefs)
  se <- sqrt(rowSums((X %*% V) * X))
  nr <- NROW(X)

  ## Calculate posterior simulation for smooths
  coefs_sim <- t(rmvn(nrep, rep(0, nrow(V)), V))
  rownames(coefs_sim) <- rownames(V)
  simDev <- X %*% coefs_sim
  absDev <- abs(sweep(simDev, 1, se, FUN = "/"))
  masd <- apply(absDev, 2, max)
  crit_s <- quantile(masd, prob = 0.95, type = 8)


  out <- list(smooth = rep(smooth, nr), by = rep(by_var, nr),
              level_1 = rep(f1, nr),
              level_2 = rep(f2, nr),
              diff = sm_diff, se = se,
              lower_s = sm_diff - crit_s * se,
              upper_s = sm_diff + crit_s*se)

  out <- new_tibble(out, nrow = NROW(X), class = "difference_smooth")
  ## Only need rows associated with one of the levels
  out <- bind_cols(out, data[r1, smooth_var])

  out
```

```
}
```

Then, `difference_smooths()` function from the file `difference_smooths.R` in the `scripts/` folder takes the output from the previous function and uses it to compute the simultaneous CI. We present the code below.

```
# this function calculates the pointwise (by calling pointwise_comparisons
   .R)
# CI and the simultaneous CI for a pairwise comparison between two smooths
difference_smooths <- function(model, smooth, n = 100, ci_level = 0.95,
    newdata = NULL,
     partial_match = TRUE, unconditional = FALSE, frequentist = FALSE, nrep
         = 10000,
     include_means = TRUE, ...) {
     if (missing(smooth)) {
         stop("Must specify a smooth to difference via 'smooth'.")
     }

     # smooths in model
     S <- gratia::smooths(model)  # vector of smooth labels - 's(x)'
     # select smooths
     select <- gratia:::check_user_select_smooths(smooths = S, select =
         smooth, partial_match = partial_match)

     sm_ids <- which(select)
     smooths <- gratia::get_smooths_by_id(model, sm_ids)
     sm_data <- map(sm_ids, gratia:::smooth_data, model = model, n = n,
         include_all = TRUE)
     sm_data <- bind_rows(sm_data)
     by_var <- by_variable(smooths[[1L]])
     smooth_var <- gratia:::smooth_variable(smooths[[1L]])
     pairs <- as_tibble(as.data.frame(t(combn(levels(sm_data[[by_var]]), 2)
         ), stringsAsFactor = FALSE))
     names(pairs) <- paste0("f", 1:2)

     Xp <- predict(model, newdata = sm_data, type = "lpmatrix")
     V <- gratia:::get_vcov(model, unconditional = unconditional,
         frequentist = frequentist)
     coefs <- coef(model)

     out <- pmap(pairs, difference_pointwise, smooth = smooth, by_var = by_
         var, smooth_var = smooth_var,
          data = sm_data, Xp = Xp, V = V, coefs = coefs, nrep = nrep)
     out <- bind_rows(out)
     crit <- qnorm((1 - ci_level)/2, lower.tail = FALSE)

     out <- add_column(out, lower = out$diff - (crit * out$se), upper = out
         $diff +
          (crit * out$se), .after = 6L)
     out
}
```

### B.7 Function for plotting limits of the pairwise comparisons

The next function has the purpose of extracting the time intervals where the simultaneous CI does not cover zero from the object where the output of `difference_smooths.R` is stored in order to overlay two rectangles that help visualize the regions where each group is statistically significant.

```
# function to obtain values for the shading regions of the pairwise
   comparison
# between the smooths

pairwise_limits <- function(dataframe) {
    # extract values where the lower limit of the ribbon is greater than
       zero
    # this is the region where the control group effect is greater

    v1 <- dataframe %>%
        filter(lower_s > 0) %>%
        select(Day)
    # get day initial value
    init1 = v1$Day[[1]]
    # get day final value
    final1 = v1$Day[[nrow(v1)]]
    # extract values where the value of the upper limit of the ribbon is
       lower
    # than zero this corresponds to the region where the treatment group
       effect
    # is greater
    v2 <- dataframe %>%
        filter(upper_s < 0) %>%
        select(Day)

    init2 = v2$Day[[1]]
    final2 = v2$Day[[nrow(v2)]]
    # store values
    my_list <- list(init1 = init1, final1 = final1, init2 = init2, final2
       = final2)
    return(my_list)

}
```

### B.8 GAM diagnostics function

In Appendix A we discuss the use of quantitative and graphical diagnostics to assess the goodness of fit of a GAM. The package *mgcv* has the function `gam.check` to provide such information, but its graphical diagnostics are made using base R graphics and therefore it is not straightforward to use them in conjunction with a *ggplot2* object. Therefore we create the function `gam_diagnostics` that uses the same code from `gam.check` but without the graphical output. In this way, we can use `appraise` from the package *gratia* to create a graphical output in *ggplot2* format and the quantitative information from `gam.check`.

```
# Function that uses the source code of `gam.check` to obtain the
   estimates
# without the plots. The source can be checked by typing `gam.check` in
   the
# console.
```

```
gam_diagnostics <- function(b, old.style = FALSE, type = c("deviance", "
   pearson",
    "response"), k.sample = 5000, k.rep = 200, rep = 0, level = 0.9, rl.
       col = 2,
    rep.col = "gray80", ...) {
    type <- match.arg(type)
    resid <- residuals(b, type = type)
    linpred <- if (is.matrix(b$linear.predictors) && !is.matrix(resid))
        napredict(b$na.action, b$linear.predictors[, 1]) else napredict(b$
           na.action, b$linear.predictors)

    fv <- if (inherits(b$family, "extended.family"))
        predict(b, type = "response") else fitted(b)
    if (is.matrix(fv) && !is.matrix(b$y))
        fv <- fv[, 1]
    gamm <- !(b$method %in% c("GCV", "GACV", "UBRE", "REML", "ML", "P-ML",
        "P-REML",
        "fREML"))
    if (gamm) {
        message("\n'gamm' based fit - care required with interpretation.")
        message("\nChecks based on working residuals may be misleading.")
    } else {
        message("\nMethod:", b$method, "  Optimizer:", b$optimizer)
        if (!is.null(b$outer.info)) {
            if (b$optimizer[2] %in% c("newton", "bfgs")) {
                boi <- b$outer.info
                message("\n", boi$conv, " after ", boi$iter, " iteration",
                    sep = "")
                if (boi$iter == 1)
                  message(".") else message("s.")
                message("\nGradient range [", min(boi$grad), ",", max(boi$
                   grad),
                  "]", sep = "")
                message("\n(score ", b$gcv.ubre, " & scale ", b$sig2, ")."
                   , sep = "")
                ev <- eigen(boi$hess)$values
                if (min(ev) > 0)
                  message("\nHessian positive definite, ") else message("\
                     n")
                message("eigenvalue range [", min(ev), ",", max(ev), "].\n
                   ", sep = "")
            } else {
                message("\n")
                print(b$outer.info)
            }
        } else {
            if (length(b$sp) == 0)
                message("\nModel required no smoothing parameter selection
                   ") else {
                message("\nSmoothing parameter selection converged after",
                    b$mgcv.conv$iter,
                  "iteration")
                if (b$mgcv.conv$iter > 1)
                  message("s")
```

```
                if (!b$mgcv.conv$fully.converged)
                  message(" by steepest\ndescent step failure.\n") else
                     message(".\n")
                message("The RMS", b$method, "score gradient at
                   convergence was",
                  b$mgcv.conv$rms.grad, ".\n")
                if (b$mgcv.conv$hess.pos.def)
                  message("The Hessian was positive definite.\n") else
                     message("The Hessian was not positive definite.\n")
            }
        }
        if (!is.null(b$rank)) {
            message("Model rank = ", b$rank, "/", length(b$coefficients),
               "\n")
        }
    }
    message("\n")
    kchck <- k.check(b, subsample = k.sample, n.rep = k.rep)
    if (!is.null(kchck)) {
        message("Basis dimension (k) checking results. Low p-value (k-
           index<1) may\n")
        message("indicate that k is too low, especially if edf is close to
            k'.\n\n")
        printCoefmat(kchck, digits = 3)
    }
}
```
